# Identification and characterization of substrate- and product-selective nylon hydrolases

**DOI:** 10.1101/2024.11.14.623603

**Authors:** Erin E. Drufva, John F. Cahill, Patricia M.B. Saint-Vincent, Alexis N. Williams, Vera Bocharova, Nikolas Capra, Flora Meilleur, Dana L. Carper, Célestin Bourgery, Kaito Miyazaki, Maina Yonemura, Yuki Shiraishi, Jerry M. Parks, Muchu Zhou, Isaiah T. Dishner, Jeffrey C. Foster, Stephen J. Koehler, Hannah R. Valentino, Ada Sedova, Vilmos Kertesz, Delyana P. Vasileva, Leah H. Hochanadel, C. Adrian Figg, Seiji Negoro, Dai-ichiro Kato, Serena H. Chen, Joshua K. Michener

## Abstract

Enzymes have evolved to rapidly and selectively hydrolyze diverse natural and anthropogenic polymers, but only a limited group of related enzymes have been shown to hydrolyze synthetic polyamides. In this work, we synthesized and characterized a panel of 95 diverse enzymes from the N-terminal nucleophile hydrolase superfamily with 30-50% pairwise amino acid identity. We found that nearly 40% of the enzymes had substantial nylon hydrolase activity, in many cases comparable to that of the best-characterized nylon hydrolase, NylC. There was no relationship between phylogeny and activity, nor any evidence of prior selection for nylon hydrolase activity. Several newly-identified hydrolases showed significant substrate selectivity, generating up to 20-fold higher product titers with Nylon 6,6 versus Nylon 6. Finally, we determined the crystal structure and oligomerization state of a Nylon 6,6-selective hydrolase to elucidate structural factors that could affect activity and selectivity. These new enzymes provide insights into the widespread potential for nylon hydrolase evolution and opportunities for analysis and engineering of improved hydrolases.

**Significance:** Nylons are common industrial polyamides with few recycling options. As an alternative to mechanical or chemical recycling, enzymes may provide a selective and energy-efficient route to deconstruct nylons from mixed waste. Several nylon hydrolases have been identified, most notably NylC, but these enzymes are all closely related and demonstrated similar activity and substrate range. In this work, we investigated a diverse set of enzymes and showed that nylon hydrolase activity is common, providing new insights into the evolution of microbial nylon hydrolysis. Unlike NylC, several enzymes demonstrated unprecedented substrate selectivity, preferentially hydrolyzing Nylon 6,6 compared to Nylon 6. These enzymes can be used to understand substrate selectivity in nylon hydrolysis and to engineer enzymes for nylon recycling from mixed waste.

## Introduction

Polyamides, commonly referred to as nylons, are synthetic polymers with remarkable versatility, a good balance of strength and ductility, and high abrasion resistance. Nylons are employed in a multitude of industries, from fashion to engineering, including the automotive and electronics sectors. Among the most common nylon variants are Nylon 6 (polyamide 6, or PA6) and Nylon 6,6 (polyamide 66, or PA66), which are distinguished by their molecular structures (Figure 1) and manufacturing processes. Since its inception, polyamide production has experienced rapid growth, with global production reaching over 8 million metric tons annually. Regrettably, polyamide manufacturing uses considerable energy inputs (197 MJ/kg) and generates substantial greenhouse gasses (10.4 kg of CO_2_ equivalent/kg) (1). These costs, combined with evolving environmental, societal, and economic demands have lent great interest and urgency to the challenge of recycling plastics such as polyamides. Unfortunately, polyamides are difficult to recycle.

**Figure 1:**
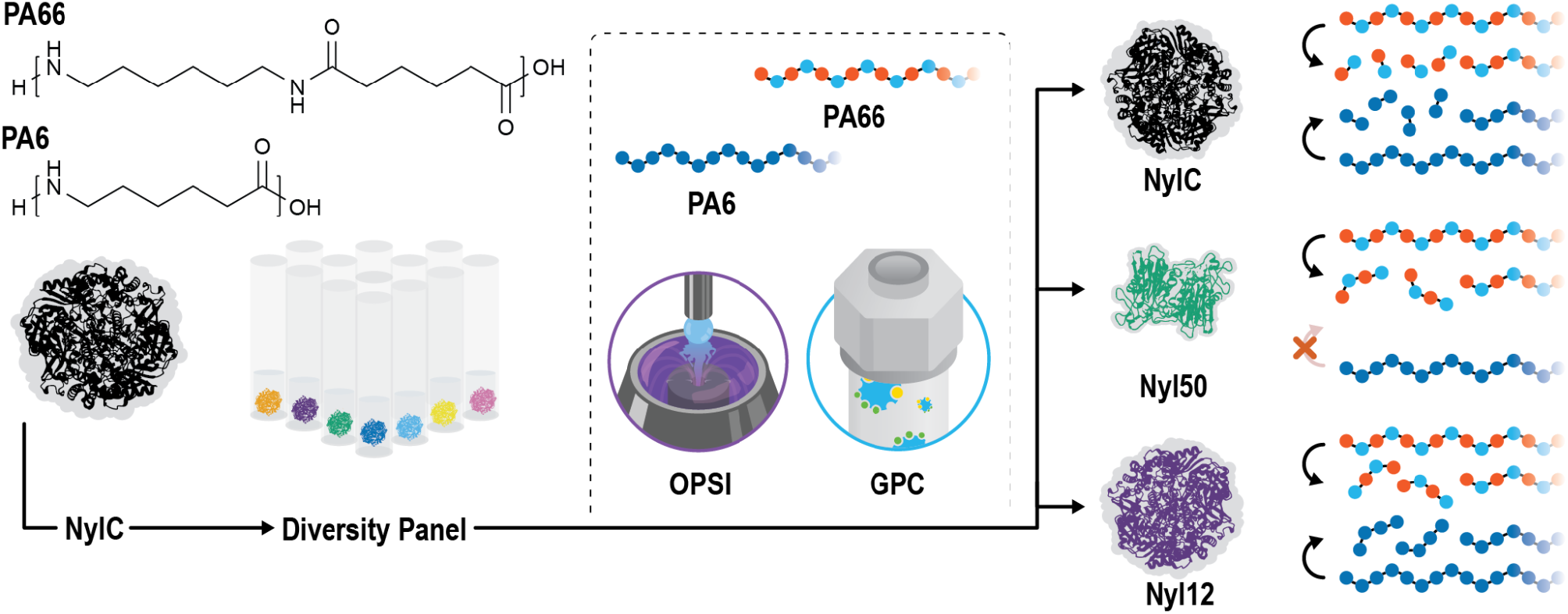
Overview of the current work. We first constructed a library of 95 homologs of NylC with 30-50% pairwise amino acid identity. We then screened the activity of the panel against PA6 and PA66 using I.DOT/OPSI-MS and GPC. We identified two new groups of enzymes, exemplified by Nyl50 and Nyl12. Both enzyme groups produced primarily tetrad products, but Nyl50 was selective for PA66 hydrolysis while Nyl12 was active on both polymers.

Nylons are often mechanically recycled, e.g. shredded and melted, to make new products. However, the process of shredding and heating alters the properties of the nylon fibers and shortens the lifespan of the materials (2–4). Chemical recycling, on the other hand, generally requires strong acid, high temperatures, produces large amounts of caustic waste, and suffers from material loss due to byproduct formation (5, 6).

Alternatively, nylon recycling via enzymatic hydrolysis and re-polymerization can be a more environmentally friendly alternative. For example, multiple studies have demonstrated the identification, characterization, and engineering of enzymes for deconstruction of polyethylene terephthalate (PET), including recent progress towards commercialization (7–11). Enzymatic PET recycling can regenerate high-quality resins from low-purity feedstocks (12). Additionally, enzymes often provide substrate and product selectivities that could generate desired hydrolysis products from mixed waste. However, polyamides are challenging substrates for enzymatic hydrolysis due to their high crystallinity.

The first nylon hydrolase, known as NylC, was discovered and characterized in a bacterium now known as *Arthrobacter* sp. KI72, which was isolated from the wastewater pond of a nylon factory. The *nylC* gene is located on a plasmid and encodes an enzyme that, in concert with NylB (6-aminohexanoate-dimer hydrolase) (13) breaks down amorphous nylon oligomers into 6-aminohexanoate monomers (14). X-ray crystallography of NylC confirmed that the enzyme autocleaves into an α and a β subunit, which then oligomerize (15). Previous mutational studies identified a quadruple mutant of NylC, termed NylC-GYAQ, with increased thermal stability and nylon-hydrolyzing activity (16, 17). Recently, several groups have explored bioprospecting for additional nylon hydrolases (18–20) or engineering NylC to improve its activity (18, 21), yielding moderate improvements in activity and thermostability.

In this work, we conducted a systematic analysis of nylon degradation by members of the N-terminal nucleophile (Ntn) hydrolase superfamily that include NylC (Figure 1). We found that nylon hydrolase activity is common in this superfamily, with comparable levels of activity to NylC but without evidence of fitness benefits from nylon hydrolysis, association of hydrolase genes with mechanisms of horizontal gene transfer, or phylogenetic association with enzyme activity. Purified proteins demonstrated a range of substrate- and product-selectivities including, to our knowledge, the first examples of enzymes that were highly selective for PA66 compared to PA6. Our analysis of the reaction products were consistent with exo- rather than endo-cleavage. Structural analysis of a PA66-selective enzyme, Nyl50, identified a potential substrate access tunnel and suggested that, unlike NylC, Nyl50 primarily assembled as a dimer in solution. Finally, we demonstrated that the oligomeric products generated by PA66 hydrolysis were readily polymerized to high molecular weight PA66. These results provide new insights into the evolution of nylon hydrolase activity, new opportunities to understand selectivity in enzymatic polymer hydrolysis, and new enzymes to engineer for efficient nylon recycling.

## Results and Discussion

### Diversity panel construction

To explore potential nylon hydrolase activity of the Ntn-hydrolase family, we searched the UniRef100 database for homologs of the query sequence, a thermostable quadruple mutant of NylC known as NylC-GYAQ. The resulting dataset of 2,839 sequences was further reduced by filtering for a minimum of 50% coverage with the query sequence and selecting the 89 sequences with the lowest maximum pairwise sequence identity (Figure S1). The dataset was supplemented with six UniProtKB sequences for a total of 95 nylon hydrolase homologs having pairwise sequence identities between 30% and 50% (Table S1).

The genes comprising this diversity panel come mostly from metagenomes so, to our knowledge, the native function of these genes has not been determined and these enzymes have not been previously characterized. Unlike NylC from *Arthrobacter* sp. K172, these genes are not near homologs of NylA (6-aminohexanoate-cyclic-dimer hydrolase) or NylB (6-aminohexanoate-dimer hydrolase). Additionally, the genes do not appear to be horizontally transferred, as differences in GC content between the genes and genomes are minimal and mobile genetic elements are not observed in the genetic neighborhood (Figure S2). Taken together, these data suggest that the native function of the genes in the diversity panel is not nylon degradation.

### Screening of diversity panel activity

Each of the genes in our diversity panel was synthesized *de novo*, cloned into pET-28a(+), and transformed into *E. coli* BL21(DE3). We then assayed the activity of each variant in crude cell lysates at 65 °C. We generated polyamide substrates for activity tests by cryomilling additive-free pellets of PA6 and PA66 to average sizes of 580 μm and 680 μm, respectively (Figure S3). The powders were then rinsed with methanol to separate small cyclic byproducts formed during polymerization. Enzymatic activity was assessed using 100 mg/mL of washed PA6 or PA66 powder or 1.25 mg/mL of the PA66 rinsate. Product formation was monitored by immediate drop-on-demand technology (Dispendix GmbH, Stuttgart, Germany) coupled with open port sampling interface-mass spectrometry (I.DOT/OPSI-MS) (22) and quantified by comparison to synthesized standards of the PA66 monomer and dimer (Figures S4 and S5).

When tested with PA66, 16/95 (15%) and 38/95 (40%) of the tested enzymes generated detectable quantities of linear monomer and dimer, respectively (Figure 2 and Figure S4). Most of the active enzymes produced primarily the linear dimer of PA66, while predominant production of the linear monomer, for example by NylC, was less common. No diacid or diamine products were observed with PA66. The extent of hydrolysis for the PA66 powder and PA66 rinsate were proportional, though much higher with the rinsate (Figure S6). We did not test for expression of the inactive members of the diversity panel, nor did we screen for activity at mesophilic temperatures, so this measure of the frequency of nylon hydrolase activity was likely an underestimate. Intriguingly, nylon hydrolase activity did not have an apparent relation to phylogeny but instead was broadly distributed across the superfamily. Structural modeling using AlphaFold2 predicted that each enzyme would adopt a fold similar to NylC. All active enzymes were modeled with high confidence, with average predicted local distance difference (pLDDT) scores >85. Although some inactive enzymes were also modeled with high confidence, many had extended low-confidence regions at their termini (Figure S7).

**Figure 2:**
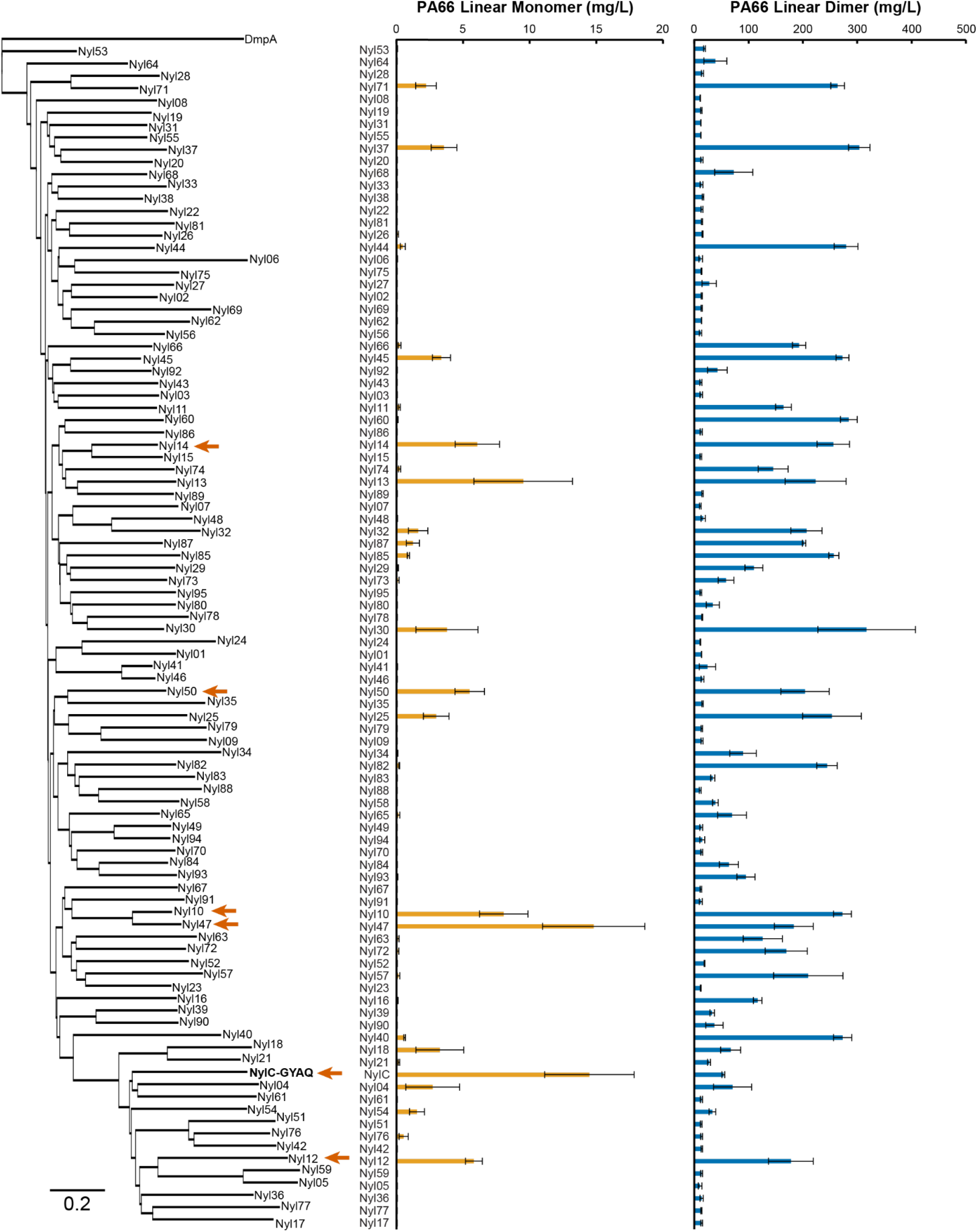
Hydrolytic activity of NylC homologs with PA66. Enzymes were expressed in *E. coli* BL21(DE3) and assayed in cell lysate. Lysates were incubated with 100 mg/mL of washed PA66 powder for 72 h at 65 °C. Products were assayed using I.DOT/OPSI-MS and quantified by calibration with synthesized standards. Error bars show the standard deviation calculated from three biological replicates. Red arrows highlight enzymes that were purified for further characterization.

Slightly fewer enzymes, 31/95 (33%), showed activity for PA6 hydrolysis and the active enzymes generally produced a broader distribution of product lengths than with PA66 (Figure S8). Similar to PA66, the majority of enzymes that hydrolyzed PA6 generated linear tetramer products. Production of smaller oligomers, such as linear dimer, was less prevalent and was largely observed in a clade centered on NylC. These findings indicated that the identification of potential nylon hydrolases through measurement of PA6 dimer production may yield false negatives. Finally, several enzymes showed differential activity with PA66 versus PA6, suggesting that they may be substrate-selective.

Based on these results, we conclude that nylon hydrolase activity is common in the Ntn-hydrolase superfamily. Diad production, i.e. either the PA66 linear monomer or PA6 linear dimer, is relatively rare though not exclusive to NylC. We propose that the key innovation in evolution of a bacterium that grows with nylon oligomers was not evolution of the oligomer hydrolase (23) but rather assembly on a mobile genetic element of that hydrolase with the accessory genes for diad hydrolysis and assimilation.

### Characterization of purified enzymes

Based on our initial screen in cell lysate, we selected six of the top performing enzymes for further characterization: Nyl10, Nyl12, Nyl14, Nyl47, Nyl50, and NylC-GYAQ. We purified each enzyme by ion exchange and size exclusion chromatography (Figure S9). Purified enzymes were assayed with washed PA66 powder at 65 °C or 75 °C and product formation was quantified by I.DOT/OPSI-MS at 24, 48, and 72 h (Figure 3A and Table S2). Based on these results, we tested four of these enzymes using washed PA6 powder at 75 °C for 72 h (Figure 3B).

**Figure 3:**
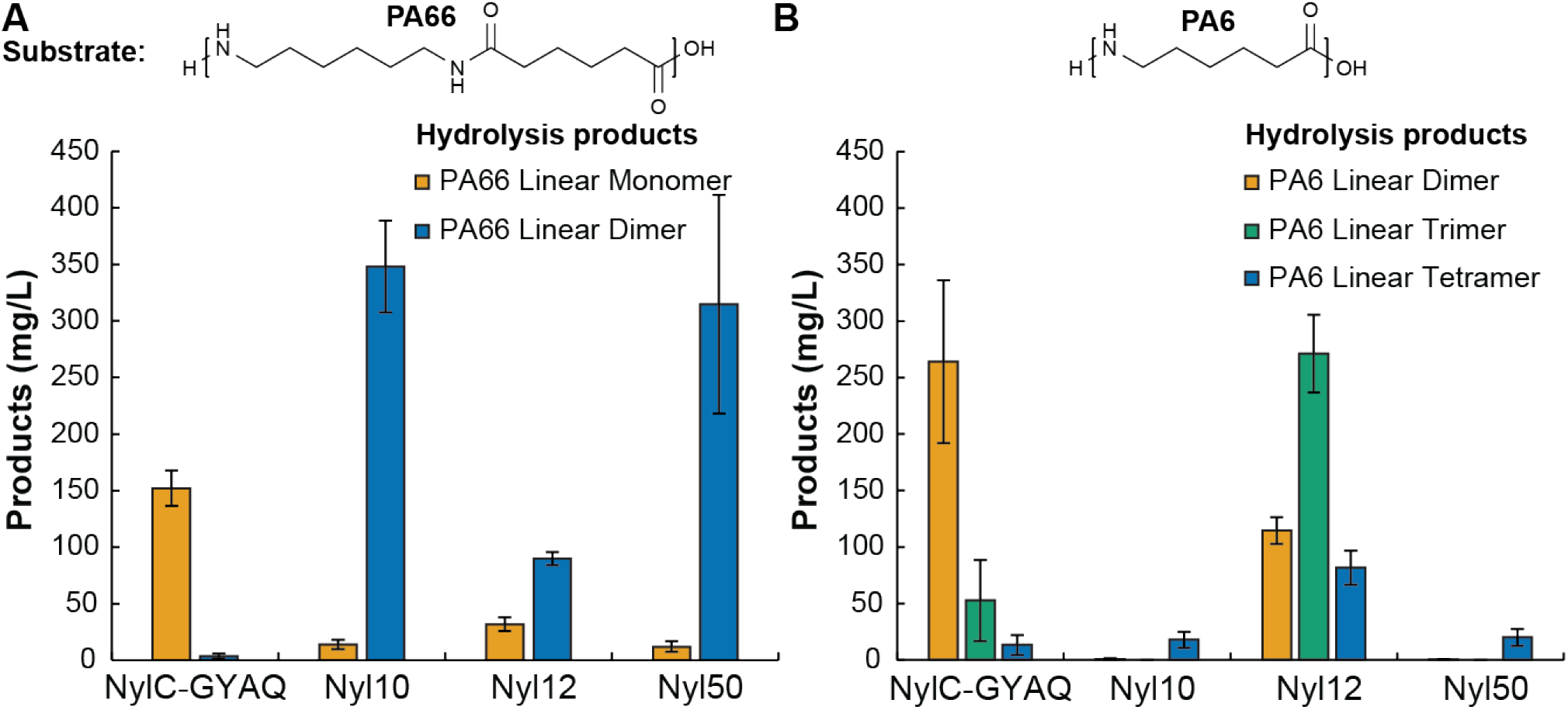
Substrate and product selectivity of top nylon hydrolases. Purified enzymes were incubated with (A) PA66 for 24 h or (B) PA6 for 72 h, both at 75 °C. Products were assayed using I.DOT/OPSI-MS and quantified by comparison to synthesized standards. Error bars show the standard deviation calculated from three biological replicates.

The top four enzymes exhibited high activity with at least one substrate. Nyl10 and Nyl50 selectively hydrolyzed PA66, generating product titers with PA66 that were 19 ± 7 and 16 ± 7-fold higher than with PA6. NylC and Nyl12 were slightly selective for PA6, generating product titers that were 2.1 ± 0.6 and 3.8 ± 0.4-fold higher than with PA66. As observed in the lysate assays, NylC was notable for primarily generating diad products: the PA6 linear dimer or the PA66 linear monomer. The other hydrolases produced longer products, such as the PA66 linear dimer. In contrast to NylC, Nyl12 produced a range of linear PA6 oligomeric products, primarily the linear trimer. Notably, we did not observe production of either PA66 triad product.

For the top four enzymes, we also measured changes in the molecular weight of the PA66 residue by gel permeation chromatography (GPC). After enzyme treatment, we saw an increase in abundance of compounds with low molecular weight (<1000 g/mol, corresponding to monomers, dimers, and cyclic compounds) and a decrease in the abundance of the bulk polymer peak (10,000-100,000 g/mol) (Figure S10). However, we did not observe the appearance of oligomers with intermediate molecular weight (1,000 - 10,000 g/mol) after enzyme treatment. In combination, these results were consistent with an exo-cleavage mechanism of hydrolysis, in which small oligomers (<1000 g/mol) were cleaved from the ends of the polymer chains.

While we detected only low levels of hydrolysis products by I.DOT/OPSI-MS after incubating Nyl50 with PA6, we could not rule out the formation of larger oligomeric products that were not readily detected by this analysis. To better assess Nyl50 substrate specificity, we also analyzed the PA6 residue by GPC after incubation with Nyl50 (Figure S11). We saw no significant changes in the polymer weight distribution, suggesting that the hydrolytic activity of Nyl50 with PA6 was very low and accurately detected by I.DOT/OPSI-MS analysis..

Extraction of the rinsate was also essential for determining the true extent of hydrolysis of the high molecular weight polymer. The PA66 rinsate was primarily composed of cyclic monomer and cyclic dimer and underwent rapid enzymatic hydrolysis to yield linear monomer or linear dimer depending on the enzyme used (Figure S12). The rinsate represented approximately 1% (w/w) of the initial mass of the powder suggesting that, unless these compounds are removed, hydrolysis activity could be dominated by the conversion of small cyclic substrates at rates largely determined by mass transfer from the bulk polymer rather than hydrolysis of the high molecular weight polymer.

We also performed a protein thermal shift assay (PTSA) on one of the PA66-selective nylon hydrolases, Nyl50, to evaluate its thermostability. The melting temperature of Nyl50 was 89 °C (Figure S13), comparable to that of the engineered thermostable NylC-GYAQ (88 °C) and NylC_K_-TS (87 °C) mutants (15, 18).

### Structural analysis of Nyl50

To better understand the structural differences between NylC and Nyl50, we crystallized Nyl50 and determined 2.2 Å (PDB ID: 9CXR) and 1.85 Å (PDB ID: 9DYS) resolution X-ray crystallographic structures at room temperature (Figure 4A and Table S3). For the sake of brevity, we refer to these two structures as Nyl50-2.2 and Nyl50-1.8, respectively. For both structures, the asymmetric unit contained two Nyl50 monomers, each containing an α and a β subunit arranged in an αββα structure corresponding to the A/D dimer previously described for NylC (15). The doughnut-shaped A/B/C/D tetramer described for NylC forms around the crystallographic two-fold axis of symmetry. Additionally, the Cα-RMSD between the conserved secondary structure elements of both Nyl50 crystal structures and the NylC and NylC-GYAQ (PDB ID 5XYG and 5Y0M, respectively) were approximately 1.5 Å, demonstrating that the enzymes all adopted similar structures despite low sequence identity. Electron density was absent for the C-terminal residues (224-226) of subunit α, indicating that autocleavage had occurred to release the β subunit N-terminal catalytic residue, Thr 227, and suggesting that the 224-226 region was disordered upon its release from the active site cleft. Spherical electron density was located 2.9 Å from Thr 227 side chain oxygen atoms in both monomers. This density was best modeled by a sodium ion. The models predicted by AlphaFold2 and AlphaFold3 agreed with both crystal structures with an all heavy-atom RMSD of 1.77 Å/1.73 Å and 0.91Å/0.97 Å, respectively (Figure S14).

**Figure 4.**
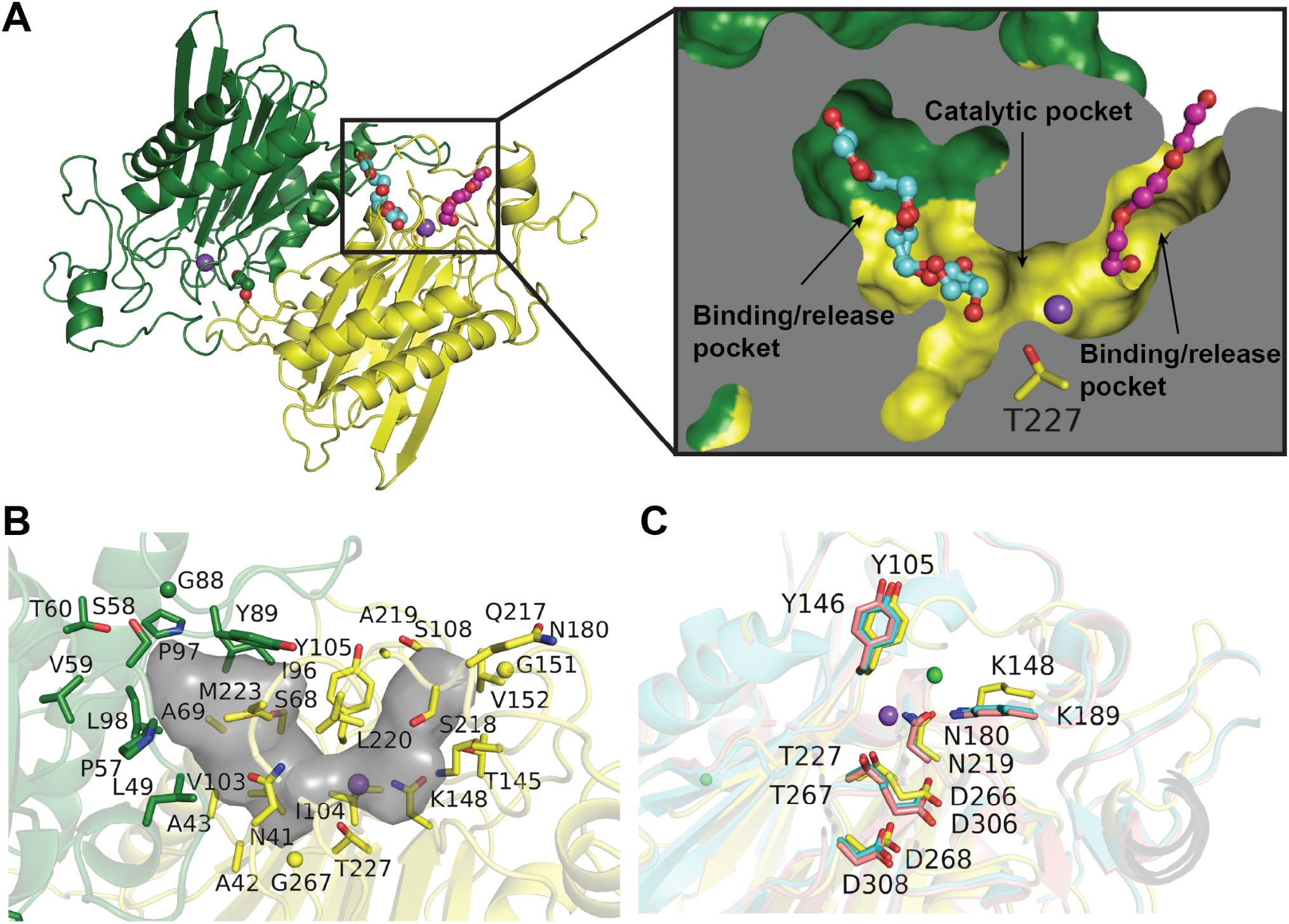
Structural analysis of Nyl50. (**A**) Crystal structure of Nyl50-1.8 showing two cleaved αβ chains in green and yellow, with the bound Na^+^ ions (purple), PG4 (cyan), EDO (green) and the modeled PGE (magenta) molecules represented as spheres. The inset shows a cut-through surface representation of the tunnel, divided into the putative pockets, with the bound PG4 molecule and the modeled PGE represented as ball-and-stick. (**B**) Residues, represented as sticks, within 5 Å of the PG4 and PGE molecules that participate in the formation of the tunnel, represented by PyVOL as a gray surface. (**C**) Comparison of the catalytic residues in the crystal structures of NylC (pink), NylC-GYAQ (cyan), and Nyl50 (yellow), shows no substantial difference in the orientation of the side chains despite the absence of ligands in NylC structures.

While the enzymes were not crystallized in the presence of a nylon-derived substrate, the presence of non-protein electron density near the active sites in both structures suggested how a polymeric substrate might access these areas. Monomer D (crystallographic molecule B) in Nyl50-2.2 contained elongated non-protein electron density. The density was observed ca. 5 Å away from Thr 227 and extended to the surface of the protein. Considering the crystallization conditions, we modeled this density as triethylene glycol (PDB ID PGE). Extra electron density was also observed at the A/D monomer interface and modeled as ethylene glycol (PDB ID EDO). Similarly, Nyl50-1.8 also contained non-protein electron density in both molecules. The electron density in monomer A was consistent with an EDO molecule. In monomer D, a well-defined, elongated non-protein electron density was observed in a position at 180º from the PGE modeled density, at about 3.5 Å from the Thr 227 sidechain. This density appeared longer than PGE and we were able to model it with tetraethylene glycol (PG4) in two alternate conformations (Figure S15).

The superposition of the two Nyl50 structures, bound to similar ligands but in different sub-sites, allowed us to identify a putative substrate access tunnel and the residues involved in tunnel formation (Figure 4A). The U-shaped tunnel extended from the surface of monomer D (Tyr 105, Leu 220) to the surface of monomer A (Ser 58, Tyr 89), leading to the catalytic residue Thr 227. Analysis with Caver (24) revealed that the tunnel entrances had radii of ∼2.5 Å while the “bottleneck” had a radius of ∼1.5 Å near the active site residue Thr 227. The tunnel-shaped cavity had an approximate volume of 631 Å^3^ that spanned both monomers. The characteristics of the tunnel satisfied the requirements for the hydrolysis of linear polyamides. A list of residues contributing to the tunnel formation is given in Table S4.

It was possible to divide the tunnel into three putative subpockets: two large subpockets for the binding/release of the substrate, one of which is located at the interface between the monomers and contains residues from both subunits, and a third smaller, catalytic pocket (Figure 4B). Structural comparison between NylC, NylC-GYAQ, and Nyl50 showed that the orientation of the side chains was nearly identical in all structures (Figure 4C), despite NylC being crystallized in its apo form. This similarity suggested that these residues were not responsible for the differing substrate and product selectivities of NylC and Nyl50, but rather that selectivity depended on details of the substrate access tunnels. However, the mechanisms of selectivity remain unclear.

### Analysis of Nyl50 oligomerization

The putative active sites of NylC-GYAQ and Nyl50 span the A/D dimer interface, suggesting that oligomerization is likely important for enzyme activity. Prior analysis of NylC and NylC-GYAQ demonstrated that tetramers were favored at moderate enzyme concentrations (0.3 mg/mL), with tetramer formation proceeding first through the formation of the A/B dimer followed by association of two A/B dimers to form an A/B/C/D tetramer (25). Consistent with this prediction, the interface between the A and B monomers in the NylC-GYAQ crystal structure (3,121Å^2^) was twice the area between the A and D monomers (1,451 Å^2^) (26).

In contrast, the Nyl50 structures showed that the A/B and A/D interfaces had similar areas, respectively 1,362 Å^2^ and 1,873 Å^2^, suggesting that Nyl50 oligomerization might favor the A/D dimer. Analytical ultracentrifugation (AUC) revealed concentration-dependent oligomerization for Nyl50, yielding tetramers at 0.15 mg/mL and dimers at 0.34 mg/mL (Figure 5A). The elution profile of Nyl50 by size-exclusion chromatography (SEC) was also consistent with dimer formation (Figure S16).

**Figure 5.**
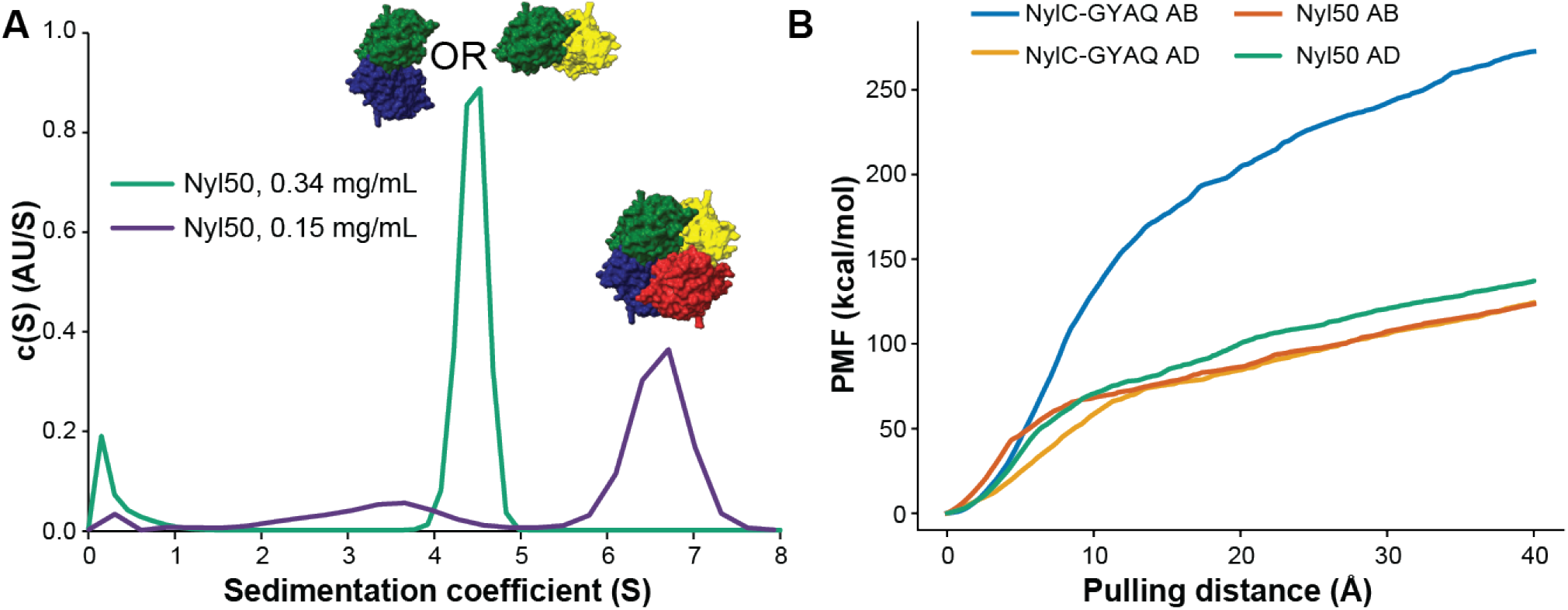
Oligomer distribution analysis of Nyl50. (**A**) c(s) distribution from analytical ultracentrifugation of Nyl50 with corresponding oligomeric state shown above each peak. (**B**) Potential of mean force (PMF) curves pulling monomer B or monomer D away from monomer A, each calculated from the exponential average work of twelve pulling pathways.

To better understand the energetic factors influencing oligomerization of NylC-GYAQ and Nyl50, we performed steered molecular dynamics simulations to estimate the strength of the AB and AD dimer interactions in NylC-GYAQ and Nyl50. We computed the potentials of mean force required to separate the two dimers of each enzyme. Consistent with interface contact areas, the Nyl50 A/D and A/B dimers required similar free energy to separate, while the A/B dimer of NylC-GYAQ required substantially more free energy to separate than the A/D dimer (Figure 5B). Combining these results, we propose that Nyl50 differs from NylC-GYAQ in primarily assembling as a dimer in solution at enzyme concentrations relevant for our assays. We speculate that Nyl50 primarily assembles as A/D dimers but cannot experimentally differentiate between A/D and A/B dimers in solution.

### Repolymerization of oligomeric products

The nylon hydrolases characterized in this work produced primarily oligomeric products. For example, Nyl50 was highly selective for PA66 linear dimer products while NylC largely made PA66 linear monomers. Typically, oligomeric products from enzymatic PA66 hydrolysis are further hydrolyzed to adipic acid and hexamethylene diamine through the action of the NylB oligomer hydrolase (27). However, PA66 oligomeric products provide potential benefits for direct repolymerization, as they retain 50-75% of the amide bonds from the original material and provide balanced stoichiometry of amine and carboxylic acid functionality. To assess the benefits of generating oligomeric hydrolysis products, we compared the polymerization of the conventional PA66 salt and chemically-synthesized PA66 linear monomer. Polymerization of the linear monomer consistently yielded PA66 with higher molecular weight than was obtained from the salt (Figure 6), suggesting that amide oligomers may be more desirable hydrolysis products than small molecule diacids and diamines.

**Figure 6.**
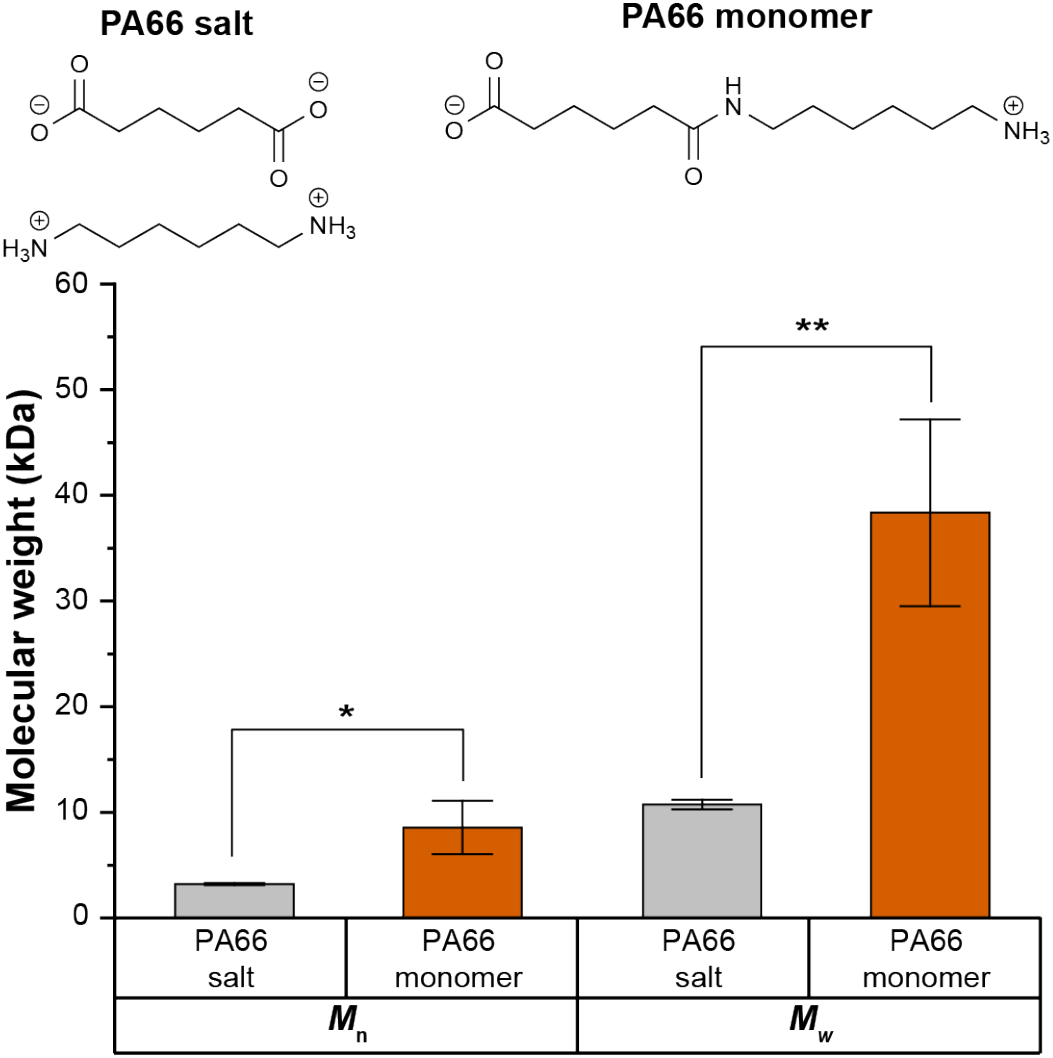
Comparison of PA66 monomer polymerization to traditional PA66 salt. Mean number-average and weight-average molecular weight (*M*_n_ and *M*_w_, respectively) values of PA66 produced from PA66 salt (gray) or PA66 monomer (orange). Polymerizations were performed in triplicate and sample means were tested for significance using a t-test (*: p < 0.05, **: p<0.01).

## Conclusions

In this work, we synthesized and characterized 95 diverse homologs of the NylC nylon hydrolase. We demonstrated that nylon hydrolase activity is common, uncorrelated with phylogeny, and most frequently generated tetrad products. Several enzymes, exemplified by Nyl50, were more active with PA66 than PA6, while other enzymes were slightly selective for PA6 or showed similar activity with both substrates. By separating soluble contaminants from the high molecular weight polymer, we demonstrated that these enzymes were capable of rapid endo-cleavage of small cyclic substrates and slower exo-cleavage of large linear polymers. Structural analysis of Nyl50 suggested that it primarily associated as a dimer in solution and identified a putative substrate tunnel for polymer access to the active site. In combination, these results provided insights into the evolution of the NylC hydrolase from a promiscuous ancestor, opportunities to understand structural factors that influence polymer substrate specificity, and starting points for engineering nylon hydrolases with enhanced activity and specificity.

## Materials and Methods

### Diversity panel construction for NylC

To explore proteins for nylon hydrolysis activity, we used the amino acid sequence of a thermostable quadruple mutant of NylC (NylC^G122/Y130/A36/Q263^, or in short NylC-GYAQ (15)), as a query for an MMseqs2-based (28) sequence search implemented in ColabFold (29) to find homologous sequences in UniRef100 (30) and acquired an alignment of 2,839 UniRef100 sequences. We then applied HHfilter (31) to calculate the number of unique homologs by filtering the alignment to a minimum of 50% coverage with NylC-GYAQ and a maximum pairwise sequence identity ranging from 25% to 95% (Figure S1), leaving 2,643 sequences. To identify 95 maximally diverse homologs for expression in a 96-well plate (one well contained NylC-GYAQ as a control), we selected the 95 sequences with the lowest maximum pairwise sequence identity (Figure S1B). Six of the 95 sequences were either fragmented, did not start with a Met, and/or containing unknown amino acids and therefore were replaced by their closest matched sequences in the UniProt Knowledgebase (UniProtKB) (32) based on BLASTP (33) searches. The resulting diversity panel consists of 89 UniRef100 sequences and six UniProtKB sequences. The pairwise sequence identities between these sequences are between 30% and 50%.

### Synthesis and Expression of Enzymes

The DNA sequences of the diversity panel enzymes and NylC were codon-optimized for *E. coli* and synthesized by GenScript (Piscataway, NJ, USA) in pET-28a(+) with XbaI/EcoRI cloning sites. Plasmids were transformed into BL21(DE3) Competent *E. coli* (New England Biolabs, Ipswich, MA, USA) according to the manufacturer’s protocol. Sequences were confirmed by Sanger sequencing (Eurofins, San Diego, USA) after transformation.

Reagents and supplies for enzyme expression, purification, and testing were purchased from VWR. For testing of the diversity panel, 96-well deep well plates containing 400 µL of Luria Bertani (LB) media per well supplemented with 50 µg/mL kanamycin were grown overnight at 37 °C with shaking. 10 µL of each preculture was transferred to a new 96-well plate containing 400 µL of terrific broth (TB) supplemented with 50 µg/mL kanamycin. The cultures were grown at 37 °C with shaking until the OD_600_ reached 0.5. Protein expression was induced by adding 0.5 mM isopropyl ß-D-1-thiogalactopyranoside (IPTG) and growing overnight at 20 °C with shaking. Plates were centrifuged at 4 °C and 3500 rpm for 30 min. The supernatant was discarded, and cells were resuspended in 200 µL of cell lysis buffer (20 mM KPO_4_ (pH 7.4), 1 mg/mL lysozyme, 1 mM phenylmethylsulfonyl fluoride (PMSF), and 10% glycerol). Cells were lysed by shaking on a platform plate shaker for 2 h at room temperature. Cell debris was removed via centrifugation at 4 °C and 3500 rpm for 1 h, and supernatants were used for enzymatic assays.

For scale-up of a single enzyme, a single colony was inoculated into 10 mL of LB and grown overnight at 37 °C with shaking. The next day, 5 mL of culture was transferred into 500 mL of TB supplemented with 50 µg/mL kanamycin. Cells were grown for 3-4 h, until the OD_600_ reached 0.7. Protein expression was then induced by adding 0.25 mM IPTG and growing overnight at 20 °C with shaking. Cells were separated by centrifugation at 4000 rpm for 30 min at 4 °C and the supernatant was discarded. The cell pellet was suspended in 25 mL 20 mM KPO_4_ (pH 7.4) and 10% glycerol (Buffer A). Cells were then sonicated for 10 min in 10 s bursts on ice. Finally, cell debris was removed via centrifugation at 17,000 rpm for 45 min at 4 °C. The supernatant was collected and used for enzyme purification.

### Purification of Nylon Hydrolases

Enzyme purification was adapted from Kakudo et al. (34). Briefly, 25 mL of 4.1 M ammonium sulfate was added dropwise to the cell lysate over a period of 5 min with slow stirring on ice. After stirring an additional 30 min on ice, precipitated proteins were collected by centrifuging (10 min at 8000 rpm and 4 °C) and discarding the supernatant. Enzymes were resuspended in Buffer A and purified on an AKTA FPLC system outfitted with a 5 mL HiTrap Q FF column (Cytiva, Marlborough, MA, USA). NylC, Nyl10, and Nyl47 were eluted by increasing the concentration of NaCl in Buffer A step-wise in 0.1 M increments, with enzymes eluting near 0.3 M NaCl. For enzymes that did not bind to the column (Nyl12, Nyl14, and Nyl50), 10K Pierce™ Protein Concentrators Polyethersulfone (PES, Thermo Scientific) were used to concentrate enzymes in Buffer A. Enzyme elution and purity was assessed via sodium dodecyl sulfate polyacrylamide gel electrophoresis (SDS-PAGE). Enzymes were purified further by size exclusion chromatography (320 ml HiPrep Sephacryl S-200 HR preparative column, Cytiva), followed by concentration (Pierce™ Protein Concentrators PES), using Buffer A. Final purified protein concentration was determined by Bradford assay and purity was assessed by SDS-PAGE (Figure S9).

For analytical ultracentrifugation, enzymes were prepared as previously mentioned up to the SEC phase. For these experiments, samples post AEX were concentrated and further purified using a Superdex® 200 Increase 10/300 GL pre-equilibrated in AUC buffer (100mM NaCl and 20mM TrisHCl, pH 7.4). Samples eluted from the SEC column as a single peak in the dimeric size range. These samples were further concentrated using 3kDa cut-off PES concentrators and loaded into the AUC without storage of the purified protein. For crystallization, samples eluted from AEX were pooled and further purified using an HiLoad Superdex® 200 26/600 pg column pre-equilibrated in a 100mM NaCl and 20mM TrisHCl, pH 7.4 buffer. Purity was assessed by SDS-PAGE.

### Substrate preparation

Polyamide pellets were subjected to ball-milling under liquid nitrogen to produce a fine powder. The characteristic particle sizes were determined using optical microscopy (Figure S3). Following milling, the powder underwent a washing process aimed at removing the low molecular weight fractions. To achieve this, methanol in excess was added to the polyamide powder, and the mixture was stirred at room temperature for 2 hours. After the initial washing, the methanol was carefully decanted, and fresh methanol was introduced for subsequent washing. This procedure was repeated three times to ensure thorough removal of the low molecular weight components.. Finally, the washed powder was dried in a vacuum oven at room temperature for a minimum of 12 hours to ensure complete solvent removal.

### Optical microscopy

An optical microscope (Olympus BX51) was used to examine the texture and determine the sizes of the PA6 and PA66 powders. The particle size estimation was performed through image analysis using AmScope software. In the calculation of the average diameter, more than 100 particles were analyzed. The particle size was calculated based on the area weighted mean diameter following equation (1).

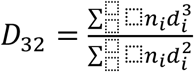

where n_i_ is the number of particles with diameter d_i,_ used to estimate the average particle size. D_32_ is the diameter of a hypothetical sphere that has the same volume-to-surface area ratio as the measured particles. The average diameter of Nylon 6 was found to be 584 µm, while for Nylon 66, it was 675 µm. The details are presented in Figure S3.

### Enzymatic Assays

For testing unpurified enzymes in high-throughput assays, 10 mg of PA6 or PA66 powder (DSM, Troy, MI, USA) was dispensed into each well of a 96-well polymerase chain reaction (PCR) plate. 75 µL 20 mM KPO_4_, pH 7.4 was added to each well along with 25 µL lysate. Plates were sealed with foil and incubated at 65 °C for 72 h unless otherwise noted. Purified enzymes were tested at a concentration of 0.3 mg/mL in the same buffer. Completed reactions were stored at -20 °C until analysis.

### I.DOT/OPSI-MS Analysis of Enzyme Activity

The immediate drop-on-demand technology (Dispendix GmbH, Stuttgart, Germany) coupled with open port sampling interface mass spectrometry (I.DOT/OPSI-MS) was used to analyze enzymatic reactions of nylons as previously described in detail (22, 35–37). Here, enzymatic reactions were diluted 1:100 (*v/v*) in high performance liquid chromatography grade water with 0.1% formic acid and 500 nM propranolol acting as a reference standard. 40 µL of diluted reactions were transferred to I.DOT S.100 96-well plates. The I.DOT system was used to eject 20 nL of sample into the OPSI-MS, which mixed the sample with a flow of 75/25/0.1 (*v/v/v*) acetonitrile/water/formic acid and transported it to the electrospray ion source of a Thermo Q-Exactive HF mass spectrometer (ThermoFisher Scientific) for accurate mass measurements and quantitation or a Sciex 7500 triple quadrupole mass spectrometer (SCIEX, Concord, Ontario, Canada) for targeted quantitation of selected linear oligomers. The Q-Exactive HF operated in positive ion mode with flow = 250 µL/min, sheath gas = 80, auxiliary gas = 40, electrospray voltage = 4 kV, mass resolution = 60,000, ion injection time = 50 ms, automatic gain control = 3e^6^, capillary temperature = 200 °C, and mass/charge (*m/z*) range = 200-2000 *m/z*. The 7500 also operated in positive ion mode with flow = 150 µL/min, gas setting 1 = 90, gas setting 2 = 60, electrospray voltage = 5.5 kV, capillary temperature = 200 °C, and dwell time = 20 ms. Multiple reaction monitoring (MRM) used the following transitions: 260.1→183.0; collision energy (CE)=26 eV (propranolol), 245.19→100.11; CE=27 eV (PA66-linear monomer), 471.35→100.11; CE=65 eV (PA66-linear dimer), 245.19→114.09; CE=26 eV (PA6-linear dimer), and 471.35→114.09; CE=65 eV (PA6-linear tetramer). Custom software (ORNL I.DOT-MS Coupler v2.50) automatically controlled I.DOT positioning, well selection, timing and droplet dispensing cycles. Additional in-house developed softwares were used for extracting data from vendor file formats, peak finding, and peak integration. For each droplet sampling event, the average mass spectrum from the resulting mass spectral peak was background subtracted and normalized to propranolol internal standard signal, correcting for droplet-to-droplet variability if present. Specific PA6 and PA66 masses were integrated over the peak to provide relative quantitation of linear and cyclic oligomers and their ion adducts (e.g., H, Na, or K). Absolute quantitation incorporated a 12-point calibration curve using standards of each linear oligomer. Dispensing throughput was 4 s/sample and each sample was measured in triplicate. Peak widths were ∼1.2 s wide. Buffer only and buffer with nylon only controls were measured for each plate.

### Gel permeation chromatography (GPC)

GPC analysis was performed on an Agilent 1260 Infinity II LC system equipped with an Agilent PL HFIPgel guard column (9 μm, 50 × 4.6 mm) and two Agilent PL HFIPgel columns (9 μm, 250 × 4.6 mm). A 0.02 M CF_3_COONa HFIP solution was used as the mobile phase at 40 °C and a flow rate of 0.3 mL min^-1^. Detection was conducted using the integrated multi-detector suite consisting of a differential refractive index (RI) detector, a multi-angle light scattering detector operating at two angles (15° and 90°), and a differential viscometer. Number-average molecular weights (*M*_n_), weight-average molecular weights (*M*_w_) and dispersities (*Ð* = *M*_w_/*M*_n_) were calculated based on calibrations with PMMA standards using the Agilent GPC/SEC software.

### Protein thermostability assays

To determine the melting temperature (T_m_) of Nyl50, two separate batches of the purified protein, expressed and purified in February 2024 and April 2024, were diluted to 1 mg/mL in Buffer A (20 mM KPO_4_ pH 7.4 with 10% glycerol). Protein thermal shift assays (PTSAs) were conducted using a QuantStudio 3 Real-Time PCR System instrument (Thermo Fisher Scientific, Waltham, MA) with a final reaction volume of 20 µL per well. Each well contained 5µL of Thermo Fisher Protein Thermal Shift Buffer (as received), 2.5 µL of Thermo Fisher Protein Thermal Shift Dye (diluted from stock solution, as received, to 8X with distilled water), and 12.5 µL of protein solution. For controls, wells containing purified protein and no dye and wells containing buffer (no protein) and dye were used. Samples were incubated on ice for 30 min, held at 25 °C for 2 min, then heated from 25–99 °C at a constant rate of 0.05 °C/s. Fluorescence was measured using the Melt Curve experiment settings in the QuantStudio Design and Analysis software and settings as described in the Thermo Fisher Protein Thermal Shift Studies user guide. Melting curves were processed and analyzed with Thermo Fisher Protein Thermal Shift software and T_m_ was determined using the derivative method in the software. No fluorescence was detected for the control samples.

### Analytical Ultracentrifugation

The AUC experiments for NylC-GYAQ and Nyl50 were performed using previously published protocols for NylC-GYAQ (25). Briefly, both enzymes were analyzed by sedimentation velocity at 20 °C in 20 mM Tris HCl, pH 7.4, and 100 mM NaCl using a Beckman-Coulter Optima XL-A analytical ultracentrifuge with the rotor pre-equilibrated at 3,000 rpm for 5 min to prevent any thermal discrepancies. Centrifugation was performed at 40,000 rpm by monitoring A280 at 5 min intervals. For sample loading, all experiments used double-sector 12-mm-thick charcoal-epon centerpieces and matched quartz windows. Experiments were analyzed using SEDFIT16.1 (38). It should be noted our buffers differed from the phosphate buffers used previously (25); however, both proteins appear to be stable in either buffer and no precipitation was observed in our experiments. This buffer discrepancy was due to ongoing structural experiments of these enzymes that were not amenable to phosphate buffers.

### Structural modeling of the diversity panel

We used the ColabFold version 1.5 (29) implementation of AlphaFold2 (39) with AlphaFold-Multimer version 3 weights (40) to model each protein in the diversity panel. MMSeqs2 (41) was used to generate multiple sequence alignments (MSAs) by searching the Uniclust _(42)_ and MGnify (43) databases, and to identify suitable templates by searching the PDB100 database (44, 45).

To identify the autocatalytic cleavage site in each protein, we generated a multiple sequence alignment with MAFFT using the L-INS-i algorithm (46). We then split each sequence into α and β chains at the site immediately preceding the residue corresponding to Thr267 in NylC. Each protein was modeled as a tetramer of heterodimers (i.e., four α and four β chains). Five models were generated and ranked according to the equation: *model confidence* = *ipTMscore·*0.8 + *pTMscore·*0.2, where *ipTMscore* is the interface predicted *TMscore* and *pTMscore* is the predicted TMscore (40). For each protein, the model with the highest score was selected for further analysis.

AlphaFold structural models and associated confidence metrics are available for download upon publication from Zenodo (https://zenodo.org link).

### Steered molecular dynamics simulations for dimer interactions

We considered two different dimer interfaces of the enzymes: one between monomer A and monomer B and the other between monomer A and monomer D. To evaluate molecular interactions at these two interfaces, we built the two dimer structures and performed nonequilibrium pulling simulations for both NylC-GYAQ and Nyl50 using a similar protocol from previous work (47). The two dimer structures of NylC-GYAQ were built from the AlphaFold2 NylC tetramer model where the four residues in each monomer (D122G, H130Y, D36A, and E263Q) were mutated by psfgen in VMD (48). The dimer structures of Nyl50 were extracted from an AlphaFold3 Nyl50 tetramer model directly (49). The AlphaFold3 model agrees with the X-ray crystal structures Nyl50-2.2 and Nyl50-1.8 with an all heavy-atom RMSD of 0.91 Å and 0.97 Å, respectively, and includes coordinates for the unresolved α subunit C-terminal tails of each monomer (Figure S14). For each dimer structure, after solvation, neutralization, and ionization to 100 mM NaCl, we minimized the system for 20,000 steps with protein heavy atoms fixed and another 20,000 steps without any fixed atoms, followed by a 10 ns equilibration. The last protein conformation from the equilibration was used as the initial state for the pulling simulations. We oriented the protein so that its three principal axes were aligned with the x, y, and z directions, re-solvated the protein into a larger water box to accommodate the pulling pathway in the z direction, neutralized with counterions and ionized to 100 mM NaCl. We then fixed the Cα atoms of four monomer A residues in the core beta sheets (residues 73, 105, 215, and 271 for NylC-GYAQ and residues 24, 63, 231, and 271 for Nyl50) and pulled the heavy atoms of the other monomer (monomer B or monomer D) at a velocity of 1 Å/ns away from monomer A for 40 ns with a 2 fs time step. The simulations were performed using NAMD (50) in NPT ensemble at 1 atm and 338 K. The CHARMM36m force field (51) and TIP3P water model (52) were used. The nonbonded interactions were calculated with a typical cutoff distance of 12 Å, while the long-range electrostatic interactions were enumerated using the particle mesh Ewald algorithm (53). Twelve pulling trajectories were sampled for each pathway. The potential of mean force was calculated according to Jarzynski’s equality (54).

### Crystallization and structure determination

Crystallization screening of Nyl50 was conducted at the National High-Throughput Crystallization Screening Center at the Hauptman–Woodward Medical Research Institute (Buffalo, NY) (55, 56). Needle-shaped crystals grew in several conditions. The commercial reagent cocktail Hampton Research (Aliso Viejo, CA) PEGRx HT-D2 (Imidazole 0.1 M pH 6.9, Polyethylene glycol 6,000 20.0% w/v) was selected to reproduce crystals. Greiner 72-well micro-batch plates were used to set up micro-batch drops under oil. Various long, rod-like shaped crystals were grown at different protein:cocktail ratios (1:2, 1:1, 2:1). Protein solution of Nyl50 at about 11 mg/mL was mixed with the crystallization cocktail up to a final volume ranging from 2 to 4 μL and subsequently overlaid with mineral oil. Crystals grew within 1 to 3 weeks. A MiTeGen MicroRT system (Ithaca, NY) was used to mount a crystal for room temperature data collection on a Rigaku HighFlux HomeLab instrument equipped with a MicroMax-007 HF X-ray generator and Osmic VariMax optics. The diffraction patterns to 2.20 and 1.85 Å-resolution were recorded using an Eiger 4 M hybrid photon counting detector. Diffraction data were integrated using the CrysAlis Pro software suite (Rigaku Inc., The Woodlands, TX) in space group *P* 2 2_1_ 2_1_ for both the crystal structures. Diffraction data were then reduced and scaled using the Aimless (57) program from the CCP4 suite (58).

For the 2.2 Å Nyl50 structure, molecular replacement using the AlphaFold2 prediction as the search model was performed with Molrep (59) from the CCP4 suite. A search for two copies of the Nyl50 monomer in the asymmetric unit was run as suggested by a Matthews coefficient calculation. Structure refinement was conducted using *phenix*.*refine* from the Phenix suite of programs (60) and manual model rebuilding was performed using the COOT molecular graphics program (61). The crystals of the 1.85 Å Nyl50 structure, determined on a second instance, belonged to the same space group of the structure at 2.2 Å with nearly identical unit cell dimensions, allowing for the previously determined structure to be used for initial refinement and electron-density map calculations. Subsequently, a few cycles of refinement using REFMAC5 (62), alternated by model building and water placement using COOT (63), were sufficient to complete the structure. Data collection and refinement statistics are shown in Supplementary Table 3. Pocket volume was calculated using PyVOL (64). Figures were generated with PyMOL (65).

## Author Contributions

E.E.D: enzyme expression and characterization; J.F.C: I.DOT/OPSI-MS analysis of hydrolysis products; P.M.B.S-V: enzyme expression and characterization; A.N.W.: oligomerization analysis; V.B.: GPC analysis, sample preparation; N.C. and F.M.: crystallization, determination and structural analysis of Nyl50: D.L.C.: I.DOT/OPSI-MS analysis of hydrolysis products; C.B.: lysate assays with PA6; K.M., M.Y., Y.S., D-I.K., and S.N.: chemical synthesis of PA66 L2; J.M.P.: computational structural analysis; M.Z.: analysis of powder properties; I.T.D and J.C.F.: synthesis of PA66 L1 and PA66 polymerization; S.J.K and C.A.F: PA6 oligomer isolation; H.R.V: initial NylC expression and characterization studies; A.S.: thermostability analysis; V.K.: I.DOT/OPSI-MS analysis and software development; D.P.V.: assisted with biochemical characterization; L.H.H.: assisted with library preparation; S.H.C.: funding acquisition, project management, diversity panel construction, steered molecular dynamics simulations; J.K.M: conceptualization, funding acquisition, project management. All authors contributed to writing and editing the final manuscript.

## Data sharing

The script and data for the diversity panel construction are available upon publication in https://github.com/serenachen0/DivSeqs.

## Acknowledgments

Research was sponsored by the Laboratory Directed Research and Development Program at Oak Ridge National Laboratory (ORNL), which is managed by UT-Battelle, LLC under Contract No. DE-AC05-00OR22725, for the U. S. Department of Energy. We thank Philip Gray for assistance with the design of Figure 1. This work used resources of the Compute and Data Environment for Science (CADES) at ORNL. Crystallization screening at the National Crystallization Center at HWI was supported through NIH grant R24GM141256.

