## Supplemental information for "Identification and characterization of substrate- and product-selective nylon hydrolases"

[1]Biosciences Division, [2]Center for Nanophase Materials Sciences, [3]Chemical Sciences Division, [4]Neutron Scattering Division, and [5]Computational Sciences and Engineering Division, Oak Ridge National Laboratory, Oak Ridge, TN, USA 37830. [6]Department of Molecular and Structural Biochemistry, North Carolina State University, Raleigh, NC, USA 27695. [7]Department of Science, Graduate School of Science and Engineering, Kagoshima University, 1-21-35 Korimoto, Kagoshima, 890-0065, Japan. [8]Department of Chemistry and Macromolecules Innovation Institute, Virginia Tech, Blacksburg, Virginia 24061, United States. [9]Department of Applied Chemistry, Graduate School of Engineering, University of Hyogo, 2167 Shosha, Himeji, Hyogo 671-2280, Japan.

**Supplemental Methods:**

***Synthesis of Methyl 6-[[6-[[(t-butoxy)carbonyl]amino]hexyl]amino]- 6-oxohexanoate:*** A 1000 mL round bottom flask (RBF) was charged with monomethyl adipate (7.5 g, 46.8 mmol, 1 equiv.) and 450 mL dichloromethane (DCM). EDC·HCl (9.87 g, 51.5 mmol, 1.1 equiv.) was added and the RBF was stirred at r.t. for 5 min until dissolved. N-Boc-1,6-hexamethylenediamine (11.15 g, 51.5 mmol, 1.1 equiv) was then added to the reaction mixture and the reaction was stirred at room temperature (r.t.) overnight. The reaction was washed with 2×250 mL 1 M HCl, 3×250 mL saturated $NaHCO_3$ solution, and 250 mL brine. The organic layer was dried over sodium sulfate, filtered, and concentrated using a rotary evaporator. The product (11.02 g, 66%) was dried overnight in a vacuum oven and used in subsequent reactions without further purification. $^1$H NMR (400 MHz, $CDCl_3$) δ 5.71 (s, 1H), 4.55 (s, 1H), 3.66 (s, 3H), 3.22 (td, *J* = 7.0, 5.8 Hz, 2H), 3.10 (q, *J* = 6.7 Hz, 2H), 2.39 – 2.25 (m, 2H), 2.18 (dq, *J* = 6.7, 3.5 Hz, 2H), 1.84 – 1.57 (m, 5H), 1.43 (s, 13H), 1.32 (p, *J* = 3.9 Hz, 4H). $^{13}$C NMR (101 MHz, $CDCl_3$) δ 174.14, 172.61, 156.23, 79.18, 51.70, 40.27, 39.22, 36.44, 33.81, 30.11, 29.56, 28.54, 26.30, 26.15, 25.26, 24.54.

***Synthesis of 6-[[6-[[(t-butoxy)carbonyl]amino]hexyl]amino]-6-oxohexanoic acid:*** To a solution of methyl 6-[[6-[[(1,1-dimethylethoxy)carbonyl]amino]hexyl]amino]- 6-oxohexanoate (11 g, 30.6 mmol) dissolved in tetrahydrofuran (THF) (110 mL) was added 1 M LiOH (110 mL). The reaction was stirred for 3.5 h and subsequently acidified to pH 4 with 1 M HCl. This solution was concentrated via rotary evaporation, and the white precipitate was filtered to provide the corresponding carboxylic acid in quantitative yield. $^1$H NMR (400 MHz, DMSO) δ 12.01 (s, 1H), 7.75 (t, *J* = 5.7 Hz, 1H), 6.77 (t, *J* = 5.7 Hz, 1H), 3.00 (q, *J* = 6.6 Hz, 2H), 2.88 (q, *J* = 6.6 Hz, 2H), 2.19 (t, *J* = 6.9 Hz, 2H), 2.04 (t, *J* = 6.8 Hz, 2H), 1.57 – 1.41 (m, 4H), 1.37 (s, 13H), 1.23 (dt, *J* = 7.7, 3.8 Hz, 4H). $^{13}$C NMR (101 MHz, DMSO) δ 174.92, 172.12, 156.04, 77.75, 40.16, 38.78, 35.58, 33.96, 29.91, 29.61, 28.74, 26.59, 26.47, 25.35, 24.63.

***Synthesis of 6-[(6-Aminohexyl)amino]-6-oxohexanoic acid trifluoroacetate salt:*** In a 500 mL RBF, 6-[[6-[[(1,1-dimethylethoxy)carbonyl]amino]hexyl]amino]-6-oxohexanoic acid (10 g, 29 mmol) was suspended in DCM (150 mL). The RBF was chilled to 0 °C in an ice bath and trifluoroacetic acid (50 mL) was added slowly to the reaction. The flask was stirred overnight, warming gradually to r.t. The majority of trifluoroacetic acid was removed by concentrating the reaction *in vacuo* and rediluting with DCM (150 mL) for 3 cycles. After the third cycle, the concentrated crude was precipitated into diethyl ether (100 mL). The ether layer was decanted, and the product was dried under vacuum to provide the product as a viscous yellow oil (8.46 g, 81%). $^1$H NMR (400 MHz, $D_2O$) δ 3.14 (t, *J* = 6.8 Hz, 2H), 2.94 (t, *J* = 7.6 Hz, 2H), 2.44 – 2.30 (m, 2H), 2.21 (td, *J* = 6.7, 4.1 Hz, 2H), 1.70 – 1.21 (m, 12H). $^{13}$C NMR (101 MHz, MeOD) δ 177.28, 175.81, 163.18, 119.66, 40.60, 40.01, 36.71, 34.56, 30.14, 28.43, 27.30, 26.95, 26.50, 25.54.

***Synthesis of 6-[(6-Aminohexyl)amino]-6-oxohexanoic acid:*** Reillex 402 (11.73 g, 112 meq) was added to a stirred solution of trifluoroacetate salt (8 g, 22.3 mmol) dissolved in deionized water (80 mL) and stirred overnight. The reaction was filtered, and the filtrate was concentrated *in vacuo* to provide a yellow residue that was further purified by triturating with ethanol. The product was isolated as a white powder (3.28 g, 60%). $^1$H NMR (400 MHz, $D_2O$) δ 3.05 (t, *J* = 6.7 Hz, 2H), 2.84 (t, *J* = 7.6 Hz, 2H), 2.11 (t, *J* = 6.8 Hz, 2H), 2.05 (t, *J* = 7.0 Hz, 2H), 1.67 – 1.31 (m, 8H),

1.31 – 1.09 (m, $J$ = 5.5, 4.1 Hz, 4H). $^{13}$C NMR (101 MHz, $D_2O$) δ 183.28, 176.64, 39.30, 38.91, 37.11, 35.54, 27.91, 26.54, 25.34, 25.33, 25.16, 25.09.

***Synthesis of Ethyl N-(t-Butoxycarbonyl)-1,6-diaminohexyl-adipate (N-Boc-66MU-OEt):*** 4-(4,6-Dimethoxy-1,3,5-triazin-2-yl)-4-methylmorpholinium chloride (DMT-MM, 1.55 g, 5.6 TCI, Japan) was added to a stirred solution of N-Boc-1,6-diaminohexane (1.12 g, 6.7 mmol, TCI, Japan) in acetone (3 mL) and Monoetyl adipate (1.2 g, 6.7 mmol, TCI, Japan) in saturated aqueous sodium bicarbonate solution (pH 8, 7 mL) at r.t. The reaction was stirred for 4 h at r.t. After removal of acetone *in vacuo*, the residual solution was extracted by EtOAc, washed with saturated aqueous $NaHCO_3$, washed with brine, dried over $Na_2SO_4$, and concentrated. The resulting crude was further purified by silica gel column chromatography (CHCl3/MeOH = 99:1) to obtain N-tBoc-66MU-OEt as clear oil (1.02 g, 2.7 mmol, 35%). $^1$H-NMR (400 MHz, $CDCl_3$) δ 5.95 (brs, 1H), 4.63 (brs, 1H), 4.07 (q, J = 7.0 Hz, 2H), 3.17 (q, J = 6.7 Hz, 2H), 3.05 (d, J = 6.0 Hz, 2H), 2.27 (t, J = 7.1 Hz, 2H), 2.14 (t, J = 6.9 Hz, 2H), 1.60 (t, J = 3.7 Hz, 4H), 1.46-1.38 (m, 13H), 1.20 (t, J = 7.0 Hz, 3H).

***Preparation of N-Boc-1,6-diaminohexyl-adipic acid (N-Boc-66MU-OH):*** A 1 M NaOH solution (7 mL) was added dropwise to a solution of N-tBoc-66MU-OEt (0.49 g, 1.3 mmol) in MeOH (7 mL) at 0 °C. The reaction was vigorously stirred for 1.5 h at 0 °C. After quenching by 1 M HCl (10 mL), the solution was extracted by EtOAc, washed with brine, dried over $Na_2SO_4$ and concentrated. Solid N-tBoc-66MU-OH obtained was subjected to the next reaction without further purification (0.39 g, 87% crude).

***Preparation of Ethyl 1,6-diaminohexyl-adipate ($H_2N$-66MU-OEt):*** TFA (10 mL) was added dropwise to a solution of N-tBoc-66MU-OEt (0.49 g, 1.3 mmol) in DCM (5 mL) at 0 °C and stirred under same temperature. After 1 h, the mixture was concentrated *in vacuo* to yield the product (0.38 g, quant. crude) as a white solid, which was subjected to the next reaction without further purification.

***Condensation of N-Boc-66MU-OH and $H_2N$-66MU-OEt:*** DMT-MM (0.45 g, 1.6 mmol) was added to the mixed solution of above prepared N-tBoc-66MU-OH in sat. aqueous $NaHCO_3$ (pH8, 10 mL) and $H_2N$-66MU-OEt in acetone (3 mL) and MeOH (10 mL) at r.t. The reaction was stirred over night at r.t. After removal of acetone and MeOH in vacuo, the residual solution was extracted by $CHCl_3$, washed with saturated aqueous $NaHCO_3$, washed with brine, dried over $Na_2SO_4$ and concentrated. Obtained crude solid was purified by silica gel column chromatography (CHCl3/MeOH = 96:4) to obtain N-tBoc-66MU-66MU-OEt as a white solid (0.20 g, 0.33 mmol, 30%). 1H-NMR (400 MHz, $CDCl_3$) δ 6.02 (brs, 2H), 5.92 (brs, 1H), 4.61 (brs, 1H), 4.10 (q, J = 7.2 Hz, 2H), 3.25-3.19 (m, 6H), 3.08 (t, J = 6.6 Hz, 2H), 2.31 (t, J = 6.9 Hz, 2H), 2.19 (s, 6H), 1.64 (t, J = 3.4 Hz, 8H), 1.50-1.42 (m, 17H), 1.32 (d, J = 3.2 Hz, 8H), 1.24 (t, J = 7.2 Hz, 3H).

***Synthesis of dimer of 66MU, 1,6-diaminohexyl-adipyl-1,6-diaminohexyl-adipic acid ($H_2N$-66MU-66MU-OH):*** A 1 M NaOH solution (10 mL) was added dropwise to the solution of above prepared N-Boc-66MU-66MU-OEt in MeOH (3 mL) at 0 °C. The reaction was vigorously stirred for 1 h at 0 °C. After quenching by 1 M HCl (15 mL), the solution was extracted by EtOAc, washed with brine, dried over $Na_2SO_4$, and concentrated. Solid N-tBoc-66MU-66MU-OH obtained was dissolved in $CH_2Cl_2$ (15 mL). To this solution TFA (15 mL) was added dropwise at 0 °C and stirred vigorously. After 1 h, the mixture was concentrated *in vacuo* and azeotroped with toluene three times to yield $H_2N$-66MU-66MU-OH (0.12 g, 0.25 mmol, 76%) as a white solid. $^1$H-NMR

(400 MHz, $D_2O$) δ 3.04-2.98 (m, 6H), 2.83 (q, J = 7.5 Hz, 2H), 2.05 (m, 8H), 1.52-1.31 (m, 18H), 1.22-1.15 (m, 6H).

***Synthesis and purification of PA6 oligomers:*** 6-Aminohexanoic acid (3 g) was weighed into a 100 mL RBF. The flask was sealed with a rubber septum and purged with argon for 15 min. The starting material was partially polymerized by heating the RBF to 220 °C for 15 min under a continuous flow of argon.

PA6 oligomers were isolated from the crude material by Preparative HPLC using a Buchi Pure C-850 FlashPrep instrument with a Prep Pure C18 column (100 Å pore size, 10 µm particle size, 250 mm length x 20 mm ID). The Nylon-6 samples were dissolved in methanol (2 mL at 50 mg/mL) and passed through a 0.2 µm PTFE syringe filter prior to injecting in the system. Using water/methanol as the A/B solvents (40 mL/min flow rate), % B increased at a rate of 2 %/min following an initial delay of 4 minutes at 0 % B. Oligomer elution was monitored by evaporative light scattering and peaks were concentrated by rotary evaporation to remove the organic cosolvent and subsequently lyophilized to remove water. The identity of each peak was confirmed by mass spectrometry (Waters Synapt G2-S mass spectrometer with a Waters Acquity I-class UPLC).

The isolated oligomers eluted as follows by the preparative HPLC method: dimer (5.0 min, MS obsv. $[M+H]^+$ = 254.2 m/z), trimer (11.6 min, MS obsv. $[M+H]^+$ = 358.3 m/z), and tetramer (15.3 min, MS obsv. $[M+H]^+$ = 471.4 m/z).

***Synthesis of PA66 salt:*** A 250 mL RBF was charged with adipic acid (12.58 g, 86 mmol) and ethanol (120 mL) and heated to 50 °C. A solution of hexamethylene diamine (10 g, 86 mmol) dissolved in 10 mL water was added to the RBF. The reaction was stirred for an additional 2 h at 50 °C and subsequently cooled to r.t. The resulting white precipitate was filtered, rinsed with additional ethanol, and dried under vacuum to provide the product in quantitative yield.

***Solid state polymerization:*** Diad or nylon salt (1 g) was weighed into a 50 mL RBF. The flask was sealed with a rubber septum and purged with argon for 15 min. The RBF was heated to 220 °C for 1 h under a continuous flow of argon. The flask was then cooled to r.t. and an aliquot of the prepolymer was collected for GPC analysis. The flask was then equipped with a flow control adapter and the prepolymer was further polymerized at 220 °C for 7 h under vacuum.

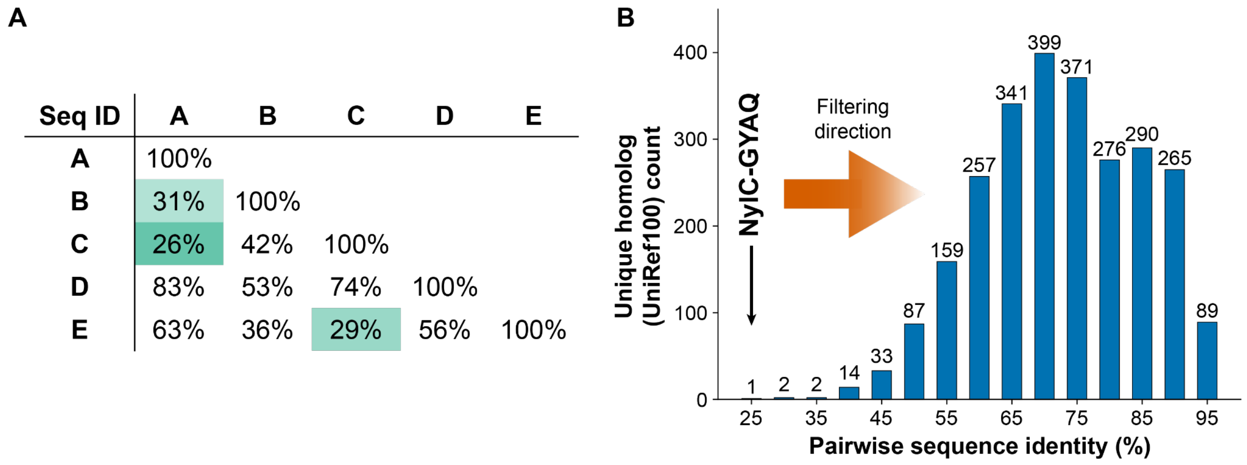


**Figure S1:** Dynamic filtering and selection of 95 maximally diverse sequences from a multiple sequence alignment of NylC-GYAQ homologs in UniRef100. (A) Hypothetical diagram describing the sequence filtering and selection order. In this example, we are selecting four sequences. We first select sequences A and C, followed by E, which all fall under the 30% pairwise sequence identity bin cutoff. We then select sequence B and complete the four-sequence set. (B) Histogram of UniRef100 unique NylC-GYAQ homologs sorted by pairwise sequence identity. The filtering criteria involve a minimum of 50% coverage with the NylC-GYAQ sequence and an increasing pairwise sequence identity cutoff from 25% to 95%, as indicated by the arrow. The number in each bin represents the number of nonredundant homologs in each identity cutoff. The only sequence in the 25% bin is the query NylC-GYAQ sequence as no other sequences satisfy the cutoff. The resulting 95 sequences share a pairwise sequence identity between 30% and 50%.

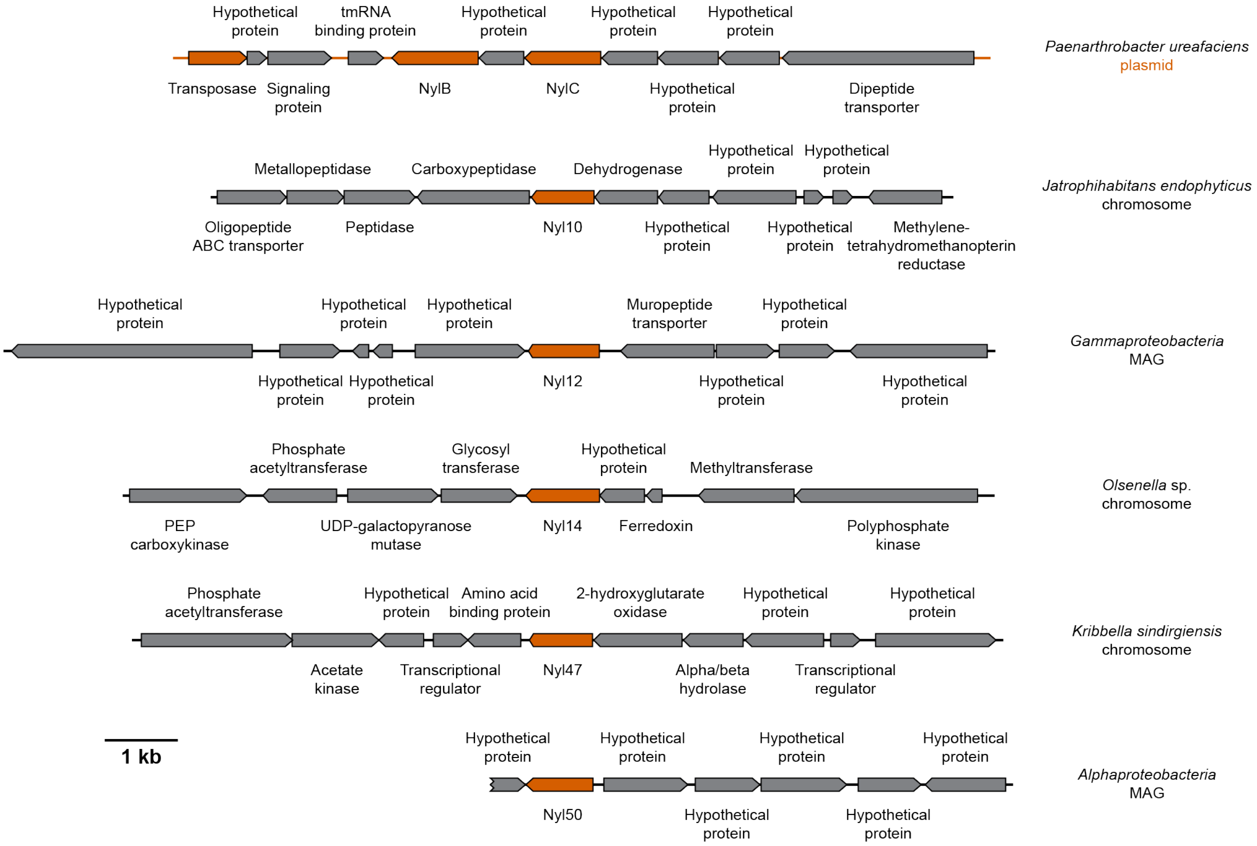


**Figure S2**: **Genomic neighborhoods of selected hydrolases.** Hydrolases are shown in the center of each row. Factors associated with horizontal gene transfer or further hydrolysis of nylon oligomers are shown in red. NylC is present on a plasmid that also contains accessory enzymes such as NylB. The other enzymes tested are chromosomal and have no such accessory enzymes in the contig or chromosome. Nyl50 is found near the end of a short contig, so little information is available on the flanking DNA.

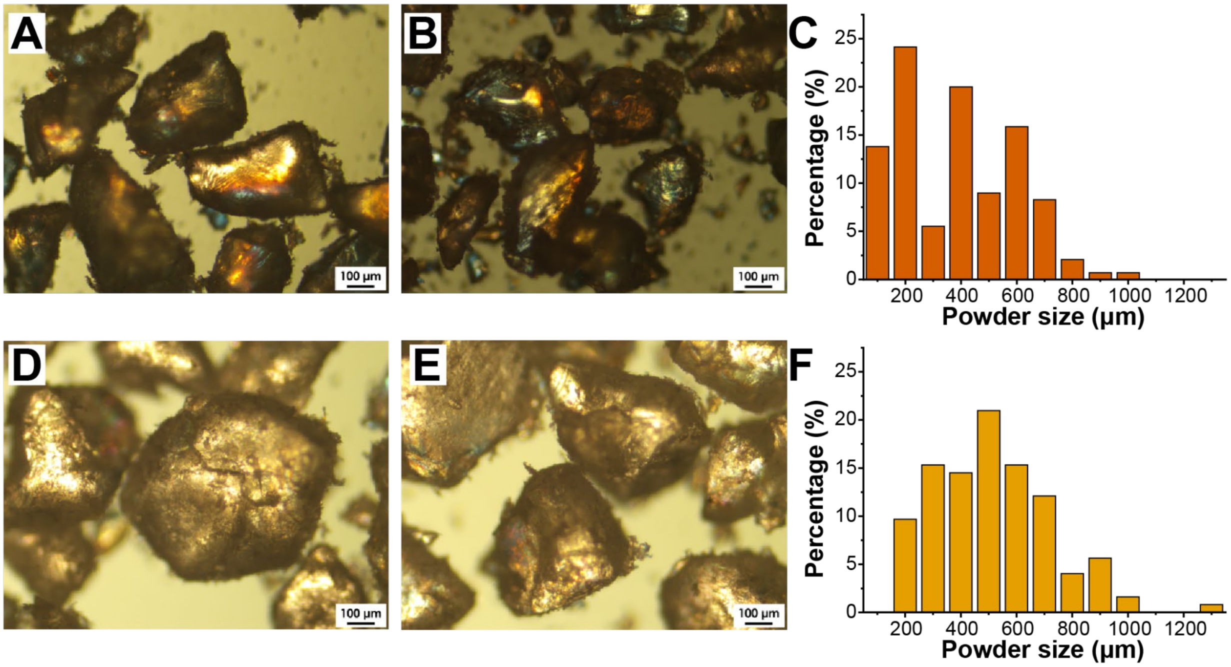


**Figure S3: Particle size distributions of PA6 and PA66 powders.** (A+B) Representative micrographs of PA6 powder. (C) Size distribution of 145 PA6 particles. The area-weighted mean diameter ($D_{3,2}$) is 580 µm. (D+E) Representative micrographs of PA66 powder. (F) Size distribution of 124 PA66 particles. The area-weighted mean diameter ($D_{3,2}$) is 680 µm.

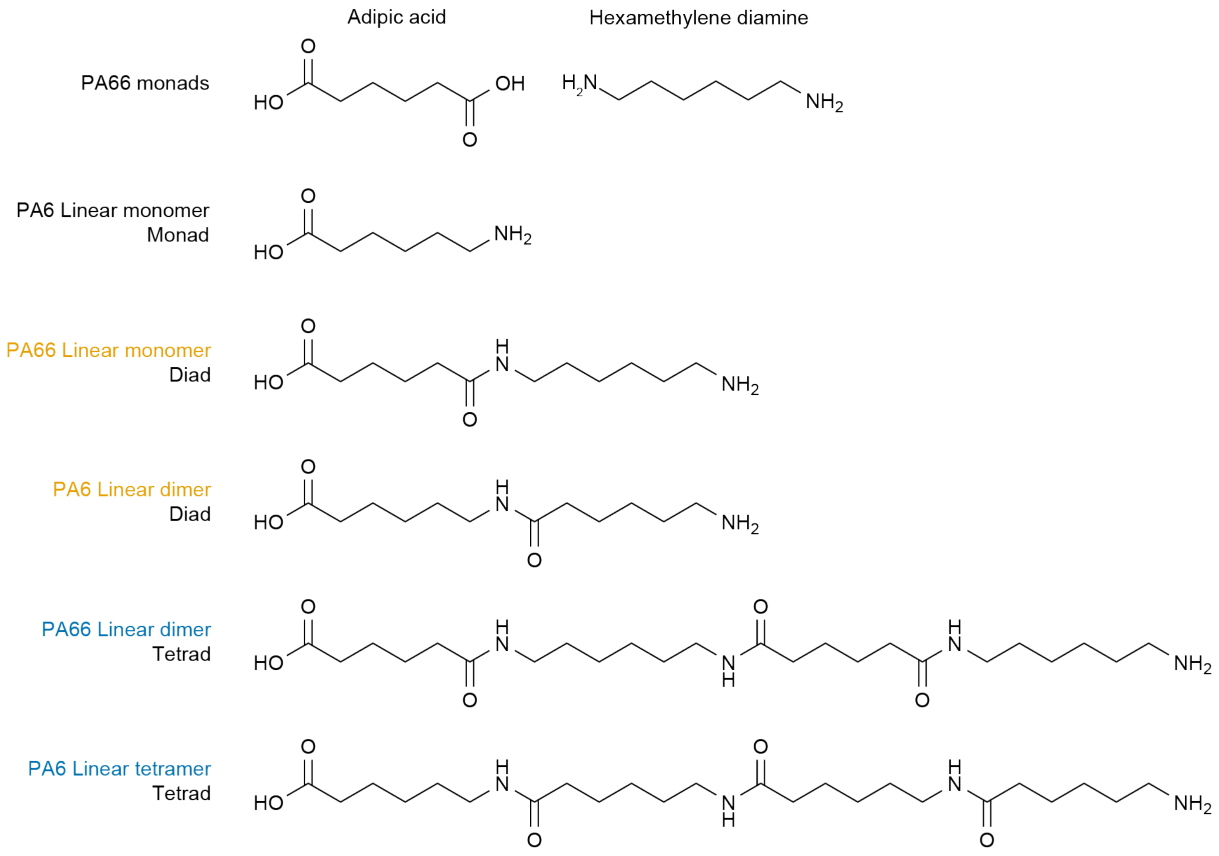


**Figure S4: Chemical structures of representative hydrolysis products.** The terms ‘monomer’ and ‘dimer’ reflect the repeat unit of the polymer, so the PA66 monomer contains one hexamethylene diamine and one adipic acid linked by one amide bond. The terms ‘diad’ and ‘tetrad’ reflect the number of amide bonds in a molecule, with a diad containing one amide bond and a tetrad containing three. Both the PA66 dimer and PA6 tetramer are tetrads.

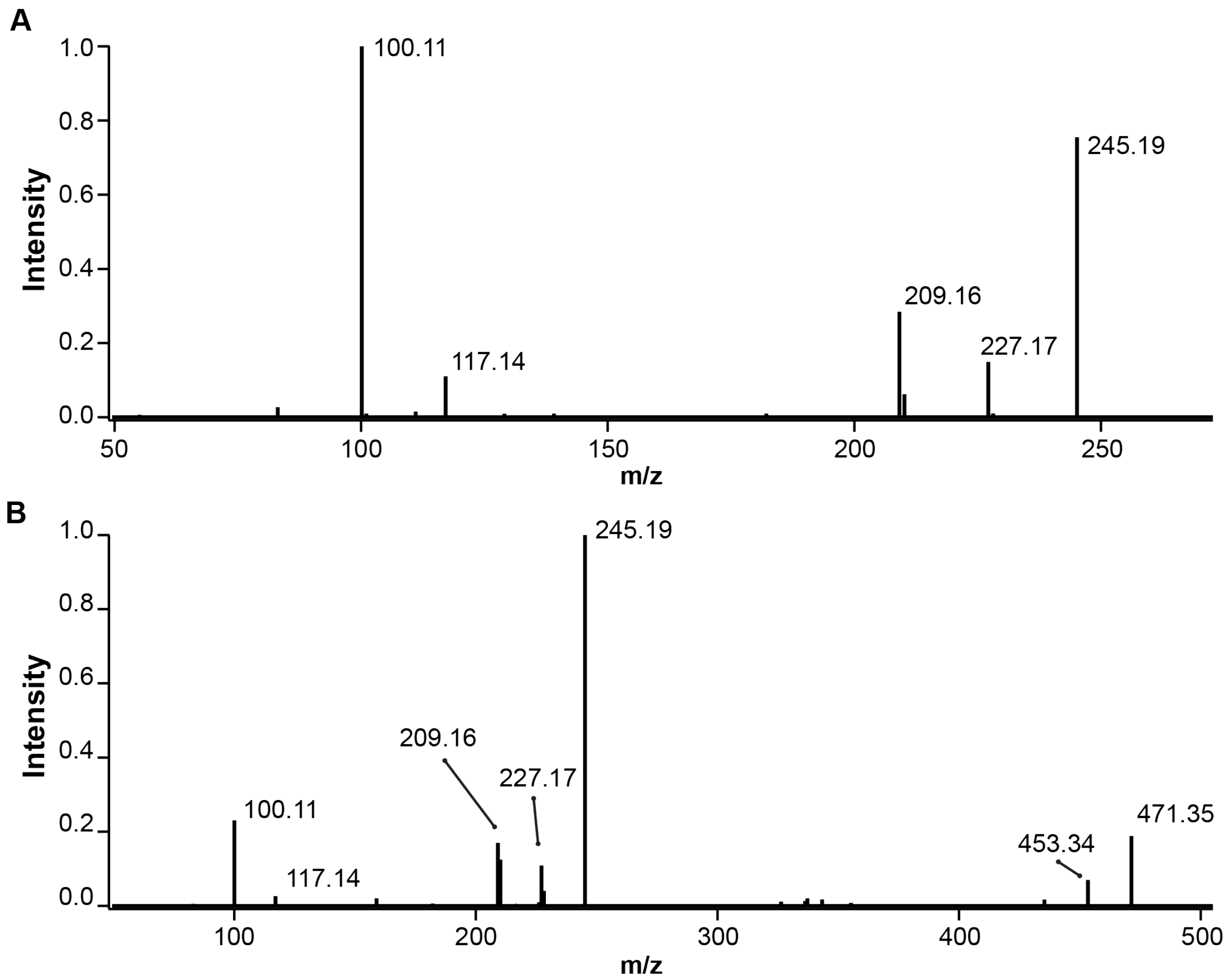


**Figure S5: Tandem mass spectra of PA66 oligomers.** (A) PA66 L1 and (B) PA66 L2. Both spectra were acquired in positive ion mode using a 35 eV collision energy.

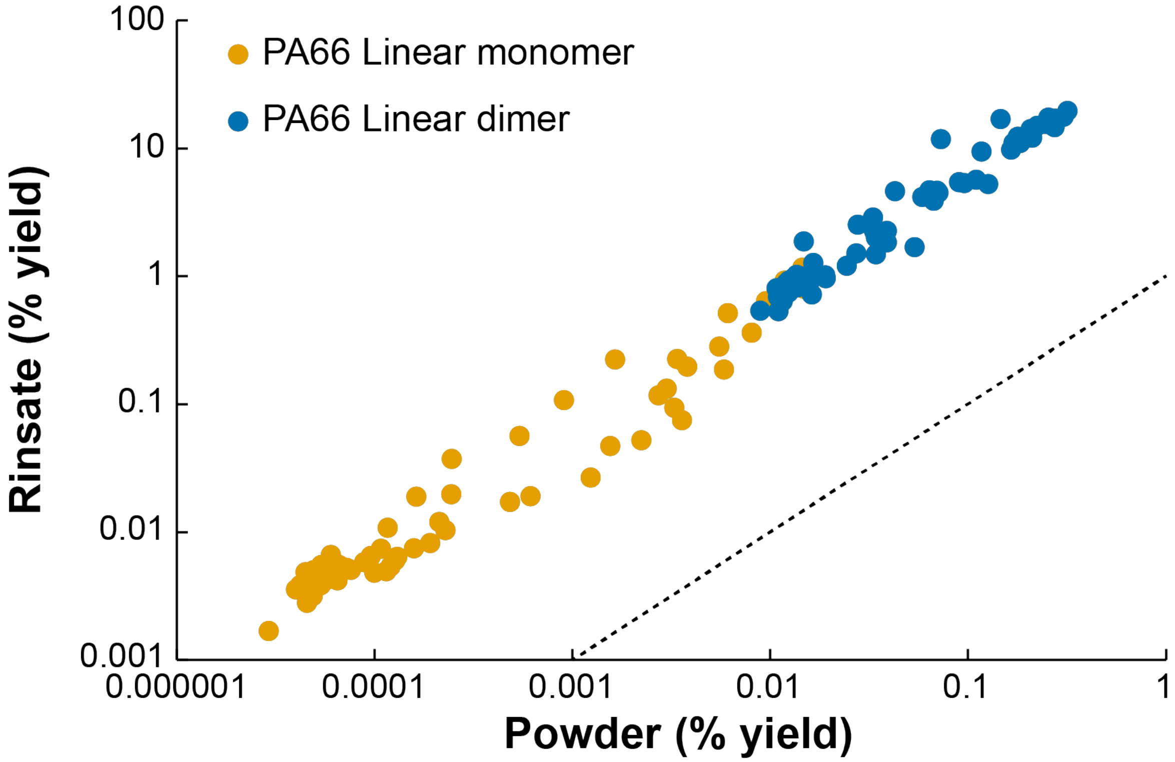


**Figure S6: Comparison of product formation with PA66 powder and rinsate.** Reactions were performed with either 100 mg/mL washed PA66 powder for 72 hours or 1.25 mg/mL PA66 rinsate for 4 hours. Each point corresponds to a single NylC homolog, as shown in Figure 2. The dashed line indicates equal yield with both substrates. Activity was largely correlated between substrates but titers were, on average, ~50% higher with rinsate than powder and yields were 120-fold higher.

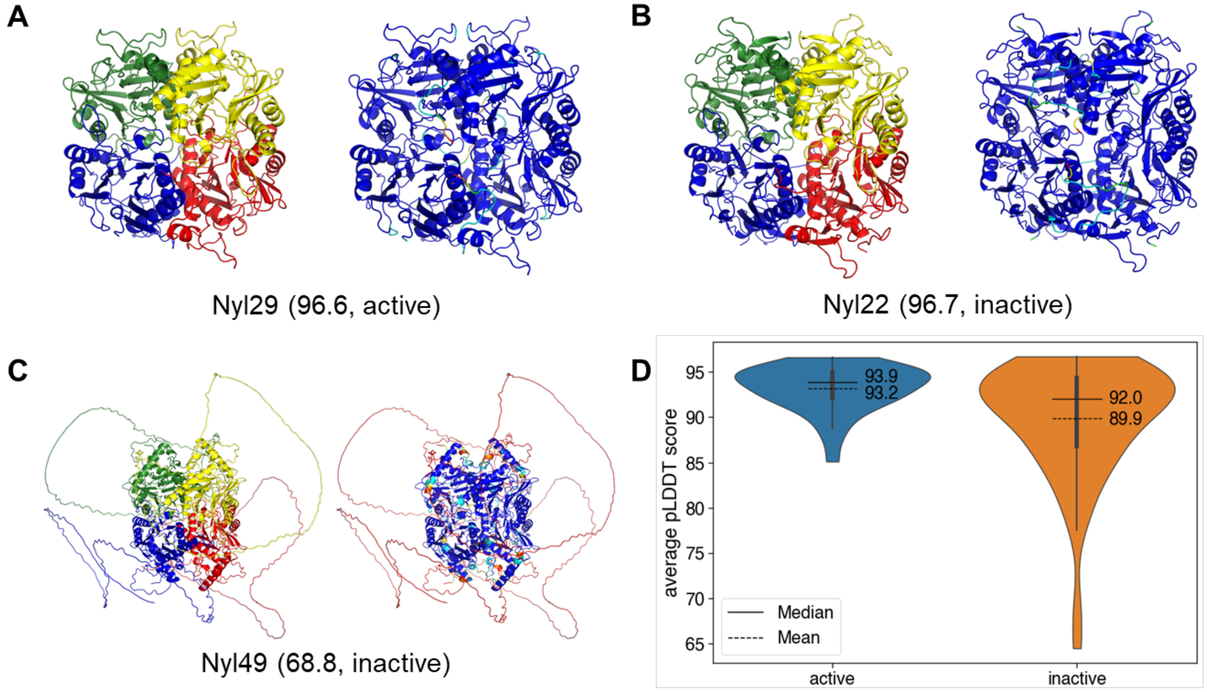


**Figure S7: Enzymatically active nylon hydrolases in the diversity panel have high AlphaFold2 confidence scores.** All active enzymes had average pLDDT scores above 85. Although some inactive enzymes were modeled with high confidence, many have extended low-confidence regions at their termini. **A**-**C**: Selected examples of a (**A**) high-confidence active, (**B**) high-confidence inactive, and (**C**) low-confidence inactive enzyme. Average pLDDT values are shown in parentheses. (**D**) Violin plot of average pLDDT scores for AlphaFold2 models of inactive and active enzymes. In **A**-**C**, each protein is colored by chain (left) and by average pLDDT score (right). Dark blue: very high confidence (pLDDT > 90), cyan: high confidence (90 > pLDDT > 70), yellow: low confidence (70 > pLDDT > 50), orange: very low confidence (pLDDT > 50).

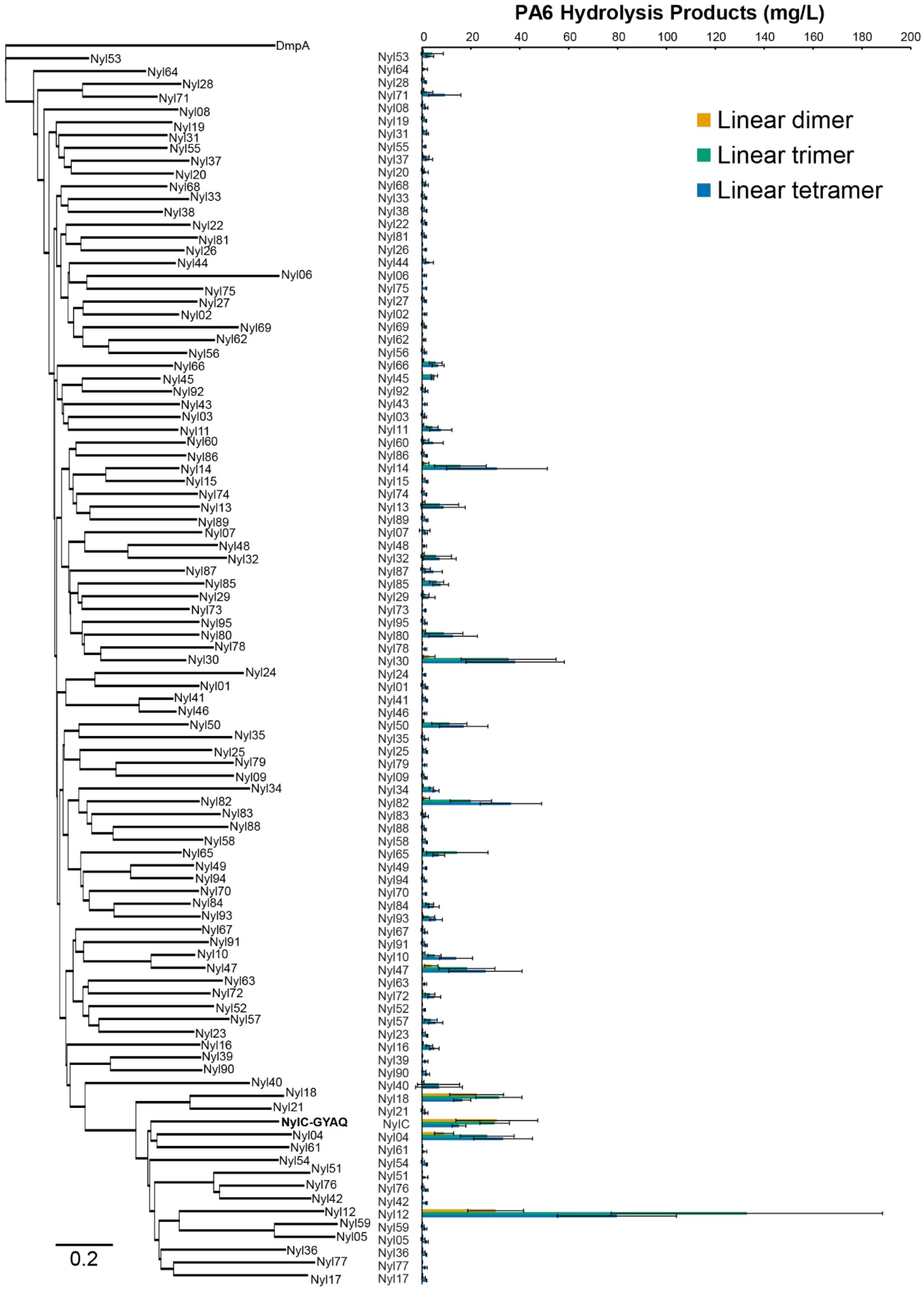


**Figure S8: Hydrolytic activity of NylC homologs with PA6.** Enzymes were expressed in *E. coli* BL21(DE3) and assayed in cell lysate. Lysates were incubated with 100 mg/mL of washed PA6 powder for 72 h at 65 °C. Products were assayed using I.DOT/OPSI-MS and quantified by comparison to synthesized standards. Error bars show the standard deviation calculated from four biological replicates.

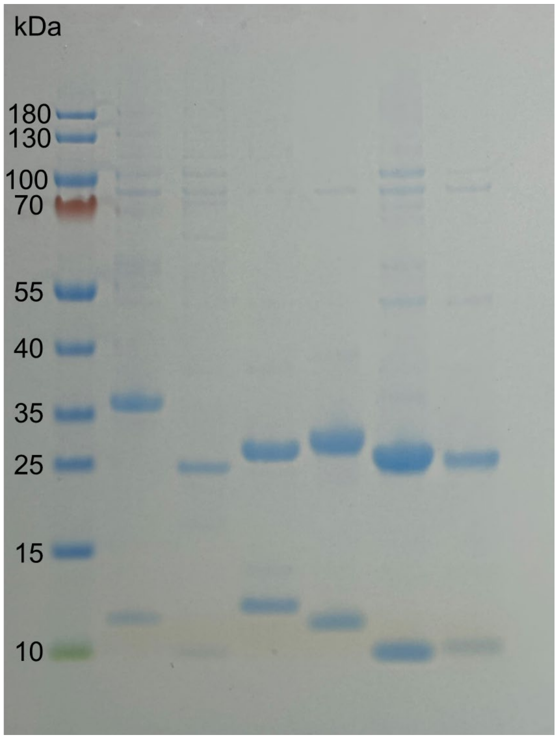

**Figure S9: Partially purified nylon hydrolases for activity assays.** From left to right: ladder, NylC-GYAQ, Nyl10, Nyl12, Nyl14, Nyl47, Nyl50.

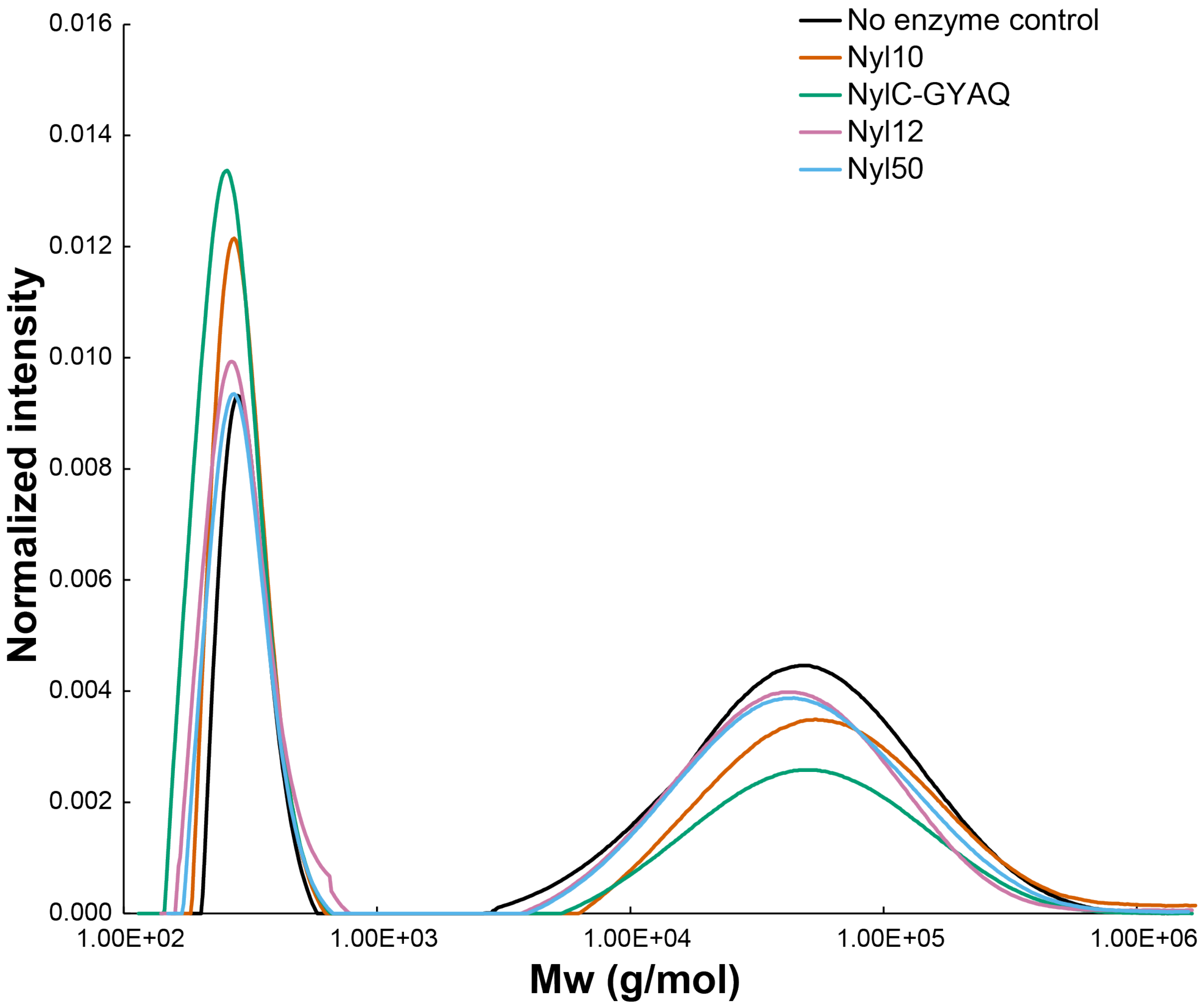


**Figure S10: GPC analysis of PA66 hydrolysis.** The area-normalized intensity is plotted against the polymer molecular weight. The samples were prepared at a concentration of 1-1.5 mg/mL in hexafluoroisopropanol. The low Mw peak is present in the control sample. We associate its presence with a certain amount of cyclic structures that were not completely removed after washing. However, the intensity and broadness of the low Mw peak changed substantially after enzymatic reactions. Quantifying these low Mw regions is challenging due to the relatively low reaction yield which brings a high variability in the position and area of the peaks. No products were observed with medium molecular weight, suggesting an enzyme-mediated exo-cleavage mechanism. For comparison, a linear PA66 10-mer would have a mass of approximately 2600 kDa.

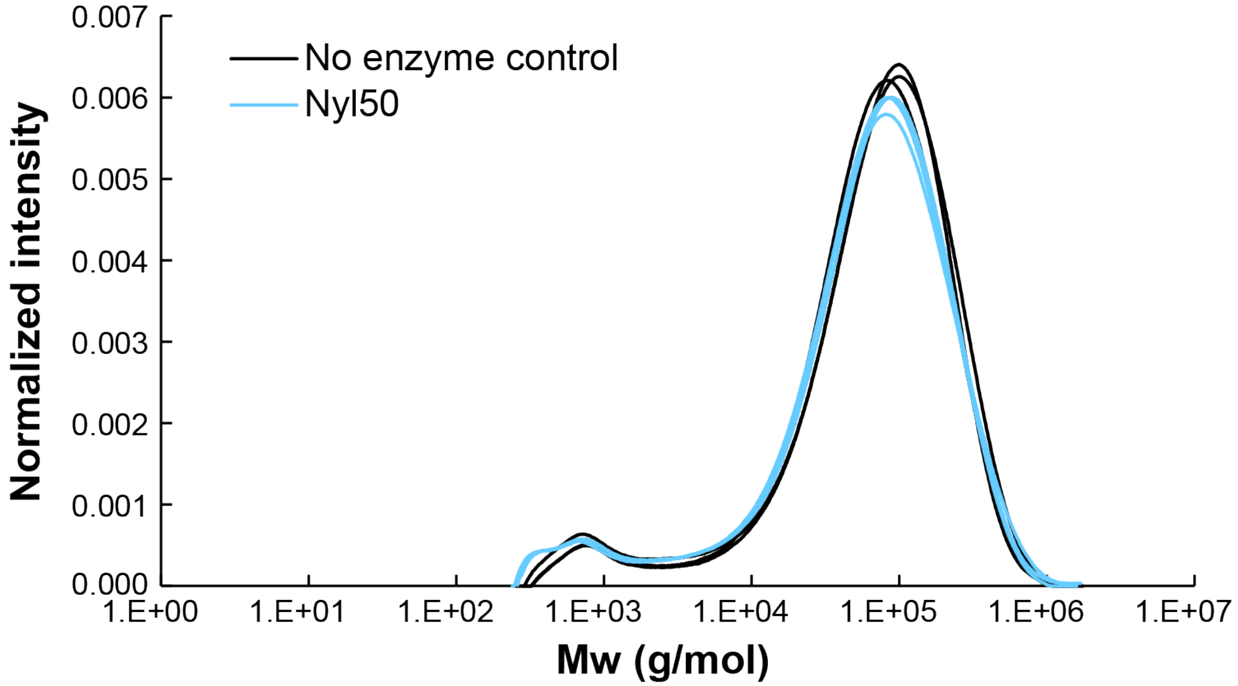


**Figure S11: GPC analysis of PA6 hydrolysis.** Analysis was conducted in triplicate as described in Figure S10, but using Nyl50 and washed PA6 powder. No products with intermediate molecular weight were identified.

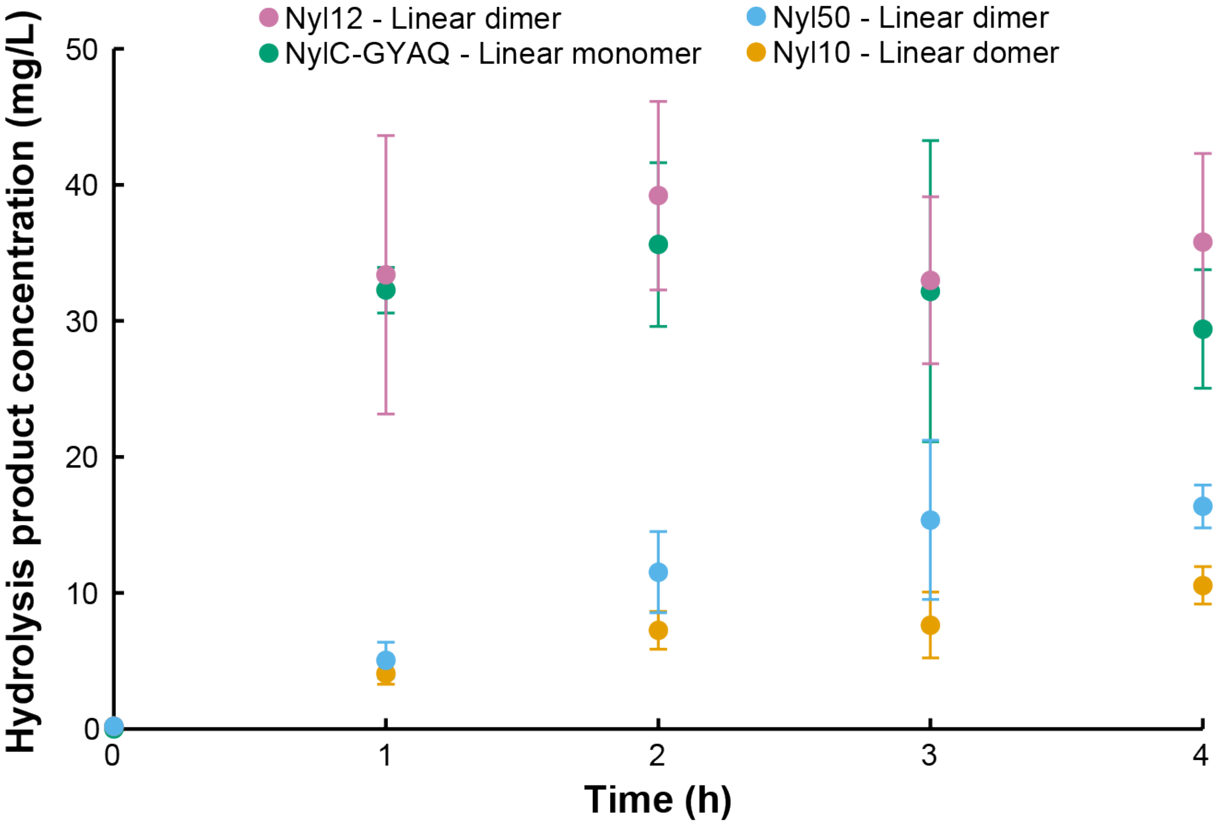


**Figure S12: Hydrolysis of PA66 rinsate.** Rinsate was incubated with 0.05 mg/mL of the indicated purified enzyme at 75 °C for 4 hours. Product formation was monitored by I.DOT/OPSI-MS and quantified by comparison to synthesized standards. Error bars show the standard deviation calculated from three biological replicates.

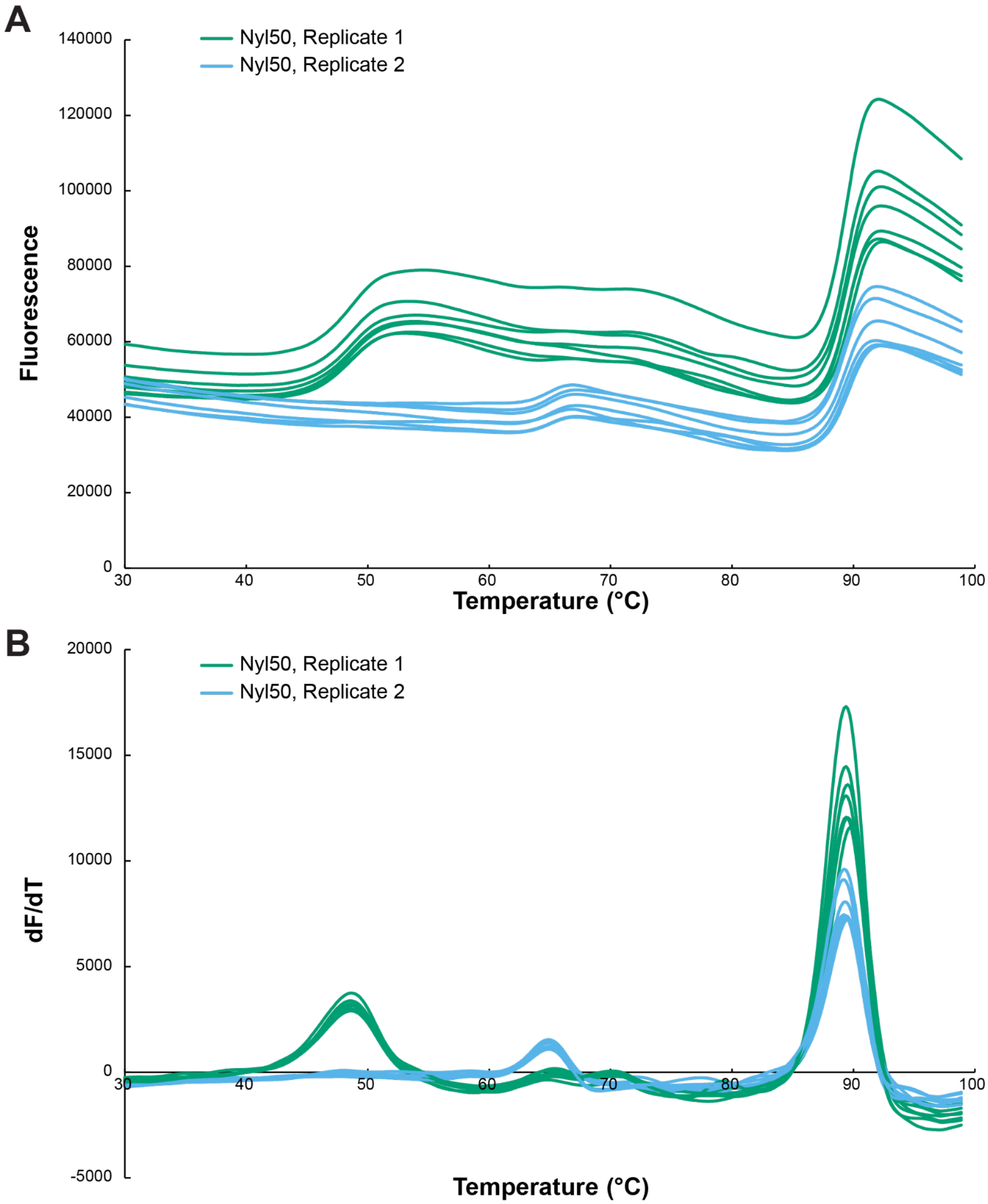


**Figure S13: PTSA melting curves for Nyl50.** Two independent preparations of Nyl50 were heated through a temperature gradient in the presence of a hydrophobic dye. Protein unfolding was monitored fluorimetrically using a quantitative thermocycler. The melting temperature was calculated with the Thermo Fisher Protein Thermal Shift Software using the derivative method to determine the inflection point of the fluorescence curve. Seven technical replicates are shown for each enzyme prep. (A) Fluorescence vs. temperature. (B) Derivative plot of the change in fluorescence (F) vs. temperature.

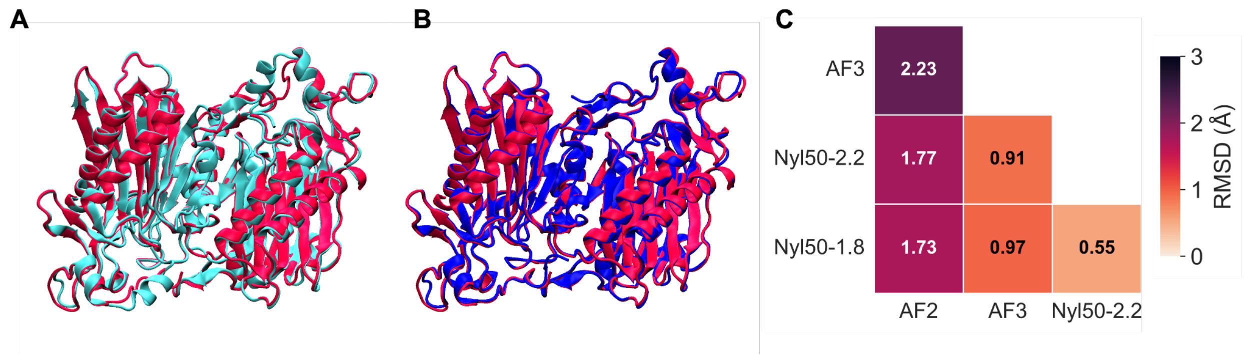


**Figure S14. Comparison of the X-ray crystallography and AlphaFold Nyl50 AD dimer structures.** (A) Aligned X-ray crystal structure Nyl50-2.2 (red) and AlphaFold2 model (cyan). (B) Aligned X-ray crystal structure Nyl50-2.2 (red) and AlphaFold3 model (blue). (C) Heat map of all heavy-atom RMSD between X-ray crystal structures and AlphaFold models.

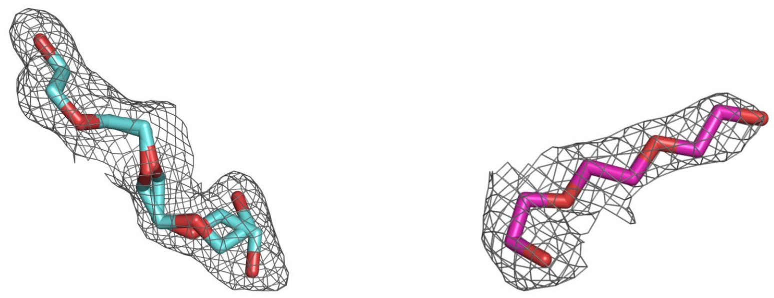

**Figure S15:** Polder maps for the tetraethylene glycol (cyan) in Nyl50-1.8 and triethylene glycol (magenta) in Nyl50-2.2 ligands, contoured at 3σ.

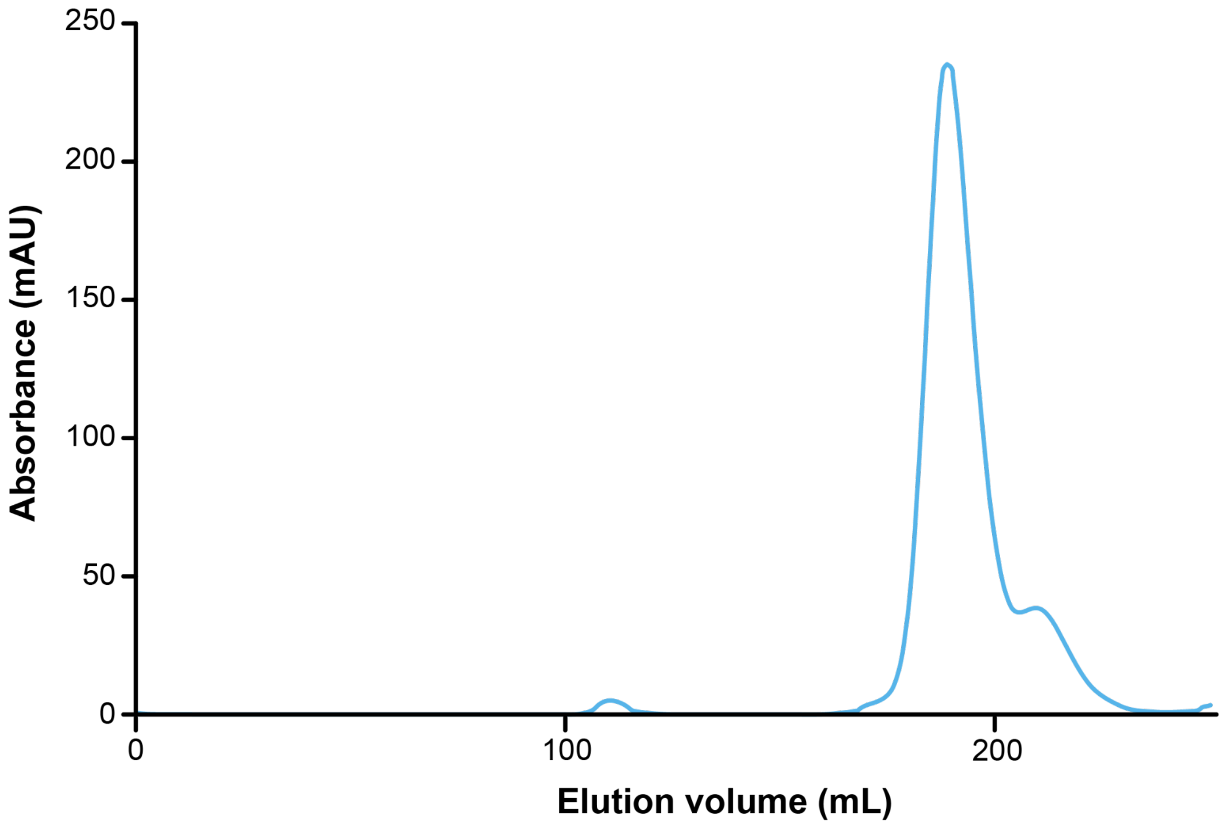


**Figure S16: Size exclusion chromatography of Nyl50.** A sharp, narrow elution peak at 189 mL corresponds to a calculated molecular weight of 74 kDa, which is consistent with a dimeric form, followed by a small peak.

**Table S2: Product titers for PA66 hydrolysis with purified enzymes**

| | Day 1 | | | | Day 2 | | | | Day 3 | | | |
|---|---|---|---|---|---|---|---|---|---|---|---|---|
| | 65 ℃ | | 75 ℃ | | 65 ℃ | | 75 ℃ | | 65 ℃ | | 75 ℃ | |
| | L1 (mg/L) | L2 (mg/L) | L1 (mg/L) | L2 (mg/L) | L1 (mg/L) | L2 (mg/L) | L1 (mg/L) | L2 (mg/L) | L1 (mg/L) | L2 (mg/L) | L1 (mg/L) | L2 (mg/L) |
| NylC-GYAQ | 110 ± 20 | 3 ± 1 | 150 ± 30 | 4 ± 2 | 150 ± 10 | 20 ± 6 | 240 ± 20 | 10 ± 2 | 190 ± 30 | 6 ± 1 | 380 ± 40 | 21 ± 9 |
| Nyl10 | 7 ± 1 | 260 ± 30 | 14 ± 4 | 350 ± 60 | 22 ± 4 | 400 ± 100 | 36 ± 4 | 440 ± 40 | 33 ± 6 | 400 ± 60 | 70 ± 10 | 550 ± 100 |
| Nyl12 | 19 ± 5 | 79 ± 1 | 32 ± 1 | 90 ± 20 | 24 ± 3 | 32 ± 4 | 56 ± 6 | 71 ± 6 | 35 ± 6 | 22 ± 6 | 67 ± 10 | 64 ± 10 |
| Nyl14 | 68 ± 9 | 106 ± 7 | 16 ± 6 | 140 ± 70 | 120 ± 10 | 90 ± 10 | 35 ± 3 | 220 ± 50 | 200 ± 40 | 110 ± 20 | 33 ± 9 | 200 ± 50 |
| Nyl47 | 18 ± 5 | 100 ± 30 | 60 ± 20 | 290 ± 120 | 30 ± 2 | 70 ± 10 | 90 ± 10 | 250 ± 30 | 60 ± 3 | 110 ± 10 | 73 ± 2 | 240 ± 40 |
| Nyl50 | 6.5 ± 0.4 | 110 ± 10 | 12 ± 2 | 310 ± 160 | 17 ± 2 | 120 ± 20 | 20 ± 5 | 300 ± 100 | 25 ± 4 | 90 ± 20 | 26 ± 3 | 350 ± 60 |

**Table S3: Data collection and refinement statistics**

| | **Nyl50_2.2** | **Nyl50_1.8** |
|---|---|---|
| **Data collection** | | |
| Wavelength (Å) | 1.54 | 1.54 |
| Space group | $P\ 2\ 2_1\ 2_1$ | $P\ 2\ 2_1\ 2_1$ |
| Unit cell dimensions | | |
| a, b, c (Å) | 54.01 96.64 105.21 | 53.88, 96.51, 105.19 |
| α, β, γ (°) | 90, 90, 90 | 90, 90, 90 |
| Resolution range (Å)* | 25.38 - 2.2 (2.28 - 2.2) | 26.73 – 1.85 (1.89-1.85) |
| Total reflections* | 125132 (12392) | 337318 (11136) |
| Unique reflections* | 28405 (2797) | 47261 (2605) |
| <I/σ>* | 10.0 (3.01) | 11.9 (2.3) |
| $CC_{(1/2)}$ * | 0.94 (0.798) | 0.996 (0.703) |
| Completeness (%)* | 98.63 (99.47) | 99.2 (90.7) |
| Wilson B-factor (Å$^2$) * | 15.4 | 22.6 |
| $R_{merge}$ | 0.177 (0.505) | 0.097 (0.662) |

**Refinement**

| | | |
|---|---|---|
| $R_{work}$/ $R_{free}$# | 0.16/ 0.20 | 0.15/0.17 |
| Number of non-H atoms in AU | 4671 | 4609 |
| protein | 4448 | 4369 |
| ligands | 18 | 32 |
| solvent | 205 | 208 |
| Average B-factor (Å$^2$) | 15.4 | 22.6 |
| RMSD | | |
| Bond lengths (Å) | 0.002 | 0.0095 |
| Bond angles (Å) | 0.49 | 1.735 |
| Ramachandran plot | | |
| % favored, outliers | 96.63, 0.17 | 97.64, 0.17 |
| Clashscore | 3.12 | 2.59 |

AU is asymmetric unit

* Values in parentheses correspond to the highest resolution shell.

### $R_{free}$ is calculated as $R_{work}$ using 5% of all reflections randomly chosen, which were excluded from structure refinement.

**Table S4: List of residues involved in tunnel formation and close contact with bound PG4.**

| Residues contributing with tunnel formation | |
|---|---|
| **Monomer A** | **Monomer D** |
| 49 LEU | 41 ASN |
| 57 PRO | 42 ALA |
| 58 SER | 43 ALA |
| 59 VAL | 68 SER |
| 60 THR | 104 ILE |
| 88 GLY | 105 TYR |
| 89 TYR | 151 GLY |
| 96 ILE | 152 VAL |
| 97 PRO | 218 SER |
| 98 LEU | 219 ALA |
| | 227 THR |
| | 267 GLY |
| **Residues in close contact with bound PG4 ( <4 Å)** | |
| 58 SER | 41 ASN |
| 89 TYR | 68 SER |
| 96 ILE | 103 VAL |
| 98 LEU | 104 ILE |
| | 227 THR |
